# A cross-cancer metastasis signature in the microRNA-mRNA axis of paired tissue samples

**DOI:** 10.1101/484816

**Authors:** Samuel C. Lee, Alistair Quinn, Thin Nguyen, Svetha Venkatesh, Thomas P. Quinn

## Abstract

In the progression of cancer, cells acquire genetic mutations that cause uncontrolled growth. Over time, the primary tumour may undergo additional mutations that allow for the cancerous cells to spread throughout the body as metastases. Since metastatic development typically results in markedly worse patient outcomes, research into the identity and function of metastasis-associated biomarkers could eventually translate into clinical diagnostics or novel therapeutics. Although the general processes underpinning metastatic progression are understood, no consistent nor clear cross-cancer biomarker profile has yet emerged. However, the literature suggests that some microRNAs (miRNAs) may play an important role in the metastatic progression of several cancer types. Using a subset of The Cancer Genome Atlas (TCGA) data, we performed an integrated analysis of mRNA and miRNA expression with paired metastatic and primary tumour samples to interrogate how the miRNA-mRNA regulatory axis influences metastatic progression. From this, we successfully built mRNA- and miRNA-specific classifiers that can discriminate pairs of metastatic and primary samples across 11 cancer types. In addition, we identified a number of miRNAs whose metastasis-associated dysregulation could predict mRNA metastasis-associated dysregulation. Among the most predictive miRNAs, we found several previously implicated in cancer progression, including miR-301b, miR-1296, and miR-423. Taken together, our results suggest that cross-cancer metastatic samples have unique biomarker signatures when compared with paired primary tumours, and that these miRNA biomarkers can be used to predict both metastatic status and mRNA expression.

## 1. Introduction

Cancer is a heterogeneous group of disorders marked by uncontrolled cell growth. It is responsible for a major financial and health burden worldwide. In the evolution of solid tissue cancers, an individual cell acquires genetic mutations that cause uncontrolled growth and division, forming a large mass, called a primary tumour. Some primary tumours acquire additional mutations that allow them to spread throughout the body as metastases. Deleterious genetic mutations disrupt normal cellular physiology by changing the structure or amount of functional molecules found within a cell. Of these molecules, RNA can be easily measured through high-throughput assays such as next generation sequencing (NGS). Since RNA is dynamically expressed within cells as they undergo the changes that lead to cancer, RNA can be used as a biomarker to potentially inform cancer diagnosis, prognosis, and treatment. RNA, especially messenger RNA (mRNA), has been used for disease prediction [14, 1, 37] and surveillance [27, 30], with promising results. MicroRNAs (miRNA), a class of small non-coding RNAs, have also been used for disease prediction [18], both individually, and in combination with mRNA [41].

MicroRNAs are often described as “master regulators” because they play an important role in maintaining gene expression [13]. As such, miRNA dysregulation may cause uncontrolled cellular proliferation through the silencing of tumour suppressors or the induction of oncogenes [13]. For example, miR-21, a so-called “oncomiR”, drives a range of cancers by inhibiting the tumour suppressor PTEN [19]. Meanwhile, some miRNAs, including members of the let-7 miRNA family, act as tumour suppressors themselves, such that their down-regulation promotes cancer [3]. In addition to their role in primary tumour development, miRNAs have been found to be associated with metastasis [20, 6]. With several miRNAs involved in vascular endothelial growth factor signalling and other key microenvironment pathways, miRNA dysregulation may induce metastases in part through promoting angiogenesis [24]. Whatever the cause, the identification of miRNA biomarkers could lead not only to the development of clinical diagnostics, but also novel therapeutics: it may be possible to develop miRNA-based chemotherapeutics by antagonizing over-expressed oncomiRs, or by replacing under-expressed microRNA tumour suppressors [8, 40, 33]. Indeed, hundreds of miRNA-based therapeutics have already been patented [8] (with Miraverson currently in Phase 2 clinical trials for the treatment of Hepatitis C [35]).

The potential diagnostic and therapeutic utility of miRNA biomarkers makes the miRNA-mRNA regulatory axis an important facet of cancer research. Yet, measuring how miRNA affects mRNA expression can be difficult because these data are collected by two different applications of NGS [17]. This results in two data sets that are often analysed independently of one another. However, when mRNA and miRNA data are collected from the same patient, and normalised appropriately, it becomes possible to integrate these data into a single analysis. Over the last decade, The Cancer Genome Atlas (TCGA) has sequenced cancerous and healthy tissue samples, from over 10,000 unique patients, representing 33 cancer types, making it the largest publicly available database of cancer transcriptomes [38]. Included in these data are 27 patients for which they performed mRNA and miRNA sequencing of a primary and a metastatic tumour sample [9]. This data set, although small, contains four expression assays per patient, providing a unique opportunity to use a paired study design to understand the role of miRNA-mRNA regulatory axis in tumour progression.

By integrating the mRNA and miRNA expression of tumour pairs, we identify a set of metastasis-associated biomarkers, enriched for canonical cancer pathways, that are capable of accurately classifying cross-cancer tissue samples as primary or metastatic. Moreover, we identify miRNAs whose metastatic biomarker profile alone can predict part of the mRNA profile too. Among the miRNAs capable of predicting cross-cancer metastatic dysregulation, we find several miRNAs previously implicated in malignancy and metastasis, including miR-301b, miR-1296, and miR-423, all of which we find correlated with several metastasis-associated genes. As targetable drivers of mRNA expression, these miRNAs could serve as good candidates for future cancer research.

## 2. Methods

### 2.1 Data acquisition

Using the R package TCGAbiolinks [10], we accessed the clinical information for all TCGA patients to look for individuals who had mRNA and miRNA count data collected for both primary and metastatic tumour samples. This query identified 27 patients across 11 unique cancers (see the Supplementary Information for patient characteristics). We then used TCGAbiolinks again to download the raw count tables for the primary and metastatic mRNA and microRNA samples for the 27 patients, yielding a total of 108 unique data sets.

### 2.2 Data normalisation and differential expression analysis

Using the 27 tissue pairs, we processed the mRNA and miRNA data in separate batches. Prior to normalisation, we removed genes with zero counts in more than half of all samples. Next, we performed effective library size normalisation of the counts with the DESeq2 package for the R programming language [2]. Using the DESeq function, we ran a differential expression (DE) analysis of the mRNA data and the miRNA data separately, comparing primary tumours with metastases via a paired sample design (by including the unique patient ID as a covariate). The normalised counts were saved for subsequent classification and regression tasks.

### 2.3 Classification of tumour metastasis

To test whether mRNA and miRNA expression could serve as biomarkers to classify tumours as primary or metastatic, we designed a machine learning pipeline with the exprso package for the R programming language [29]. At each step in the cross-validation procedure, we selected the top *{N* = 64, 128, 256}features according to the Student’s t-test, then used these top features to train a LASSO [12], random forest (RF) [21], or support vector machine (SVM) [26] model on the training set. For LASSO, each training set underwent an additional 5-fold cross-validation procedure to tune the hyper-parameter *λ*, which determines the sparsity of the LASSO model (the final *λ* was chosen to minimize the expected error across the 5-folds). Otherwise, we did not tune any hyper-parameters. We selected all features from the training sets only, making the test sets truly statistically independent.

For cross-validation, we used a modified Monte Carlo cross-validation (MCCV) procedure, whereby we randomly sub-sampled the 27 data pairs so that each training set contained 18 pairs. This design balanced the training set with regard to the tissue of origin so that this covariate would not bias the discovery of metastasis-specific diagnostic signatures. We repeated the MCCV procedure for 100 sub-samplings of the data.

### 2.4 Prediction of mRNA expression

To test whether miRNA expression alone can predict mRNA expression, we designed a regression pipeline with exprso [29]. This pipeline considered each one of the *g* = 1*…*267 differentially expressed mRNA genes as a unique outcome, treating the *m* = 1*…*569 miRNAs as predictors. To take advantage of the paired sample design, we converted the mRNA outcomes **y**_*g*_ and miRNA predictors **x**_*m*_ into new variables:

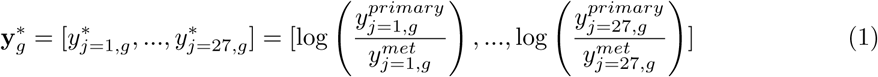

and

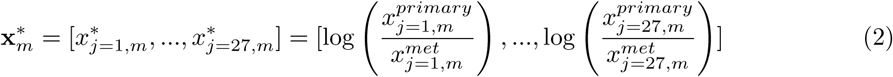

for patients *j* = 1*…*27. Thus, we fit 267 models of the form 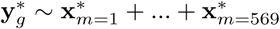. One could interpret this model as regressing the log-fold changes (LFC) of all miRNA transcripts against the LFC of one mRNA gene, allowing us to measure how well miRNA LFCs can predict mRNA LFCs.

At each step in the MCCV procedure, we randomly sub-sampled the data so that each training set contained 18 observations. Then, we selected the top *N* = 128 features according to the univariate correlation coefficient between a single miRNA and the mRNA outcome, using these top features to train an RF model as above (i.e., without hyper-parameter tuning and with all features selected from the training set).

### 2.5 Significance of regression

It can be difficult to assess the regression error of an RF model in absolute terms because the regression error depends partly on the distribution of the outcome. Even in log space, genes with higher counts can have more variance (called over-dispersion [5, 32]). For this reason, the LFCs could have higher variance too. As such, good fits of highly variable genes may have more error than bad fits of lowly variable genes, owing only to random effects.

Our objective here is to determine whether the observed root mean squared errors (RMSEs) from “Prediction of mRNA expression” are significantly better than chance. To assess this, we repeated the MCCV pipeline another 250 times, using permuted instances of the predictors instead. In other words, we randomly sampled each predictor column-wise at the start of every MCCV step. This preserves the univariate distribution of a given 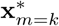, but changes the multivariate distribution for any combination of 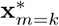 and 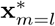. This permutation procedure gives us a *null distribution* of RMSEs for each 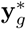. Next, we performed a one-tailed Wilcoxon Rank-Sum test to compare the distribution of observed RMSEs with the distribution of permuted RMSEs [39]. To control the false discovery rate, we adjusted the *p*-values using the Benjamini-Hochberg procedure [4].

### 2.6 Bipartite graph analysis

We trained an RF model to predict mRNA LFC based on miRNA LFC because RF tends to perform well for high-dimensional regression tasks while also allowing the analyst to interpret relative feature importance through “node purity” (see documentation for [21]). Having fit *g* = 1*…*267 models across *b* = 1*…*50 MCCV instances, we have a node purity vector which describes the importance of each miRNA *m* = 1*…*569 in fitting mRNA *g* during MCCV instance *b*:

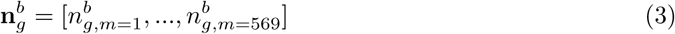

Since node purity only describes the relative importance of a feature, we convert the absolutevalues of **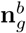** lowest rank:into a non-parametric rank vector, 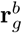, such that the smallest number receives the lowest rank

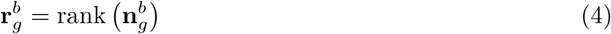

From this, we can calculate the importance of miRNA *m* for the prediction of gene *g* by

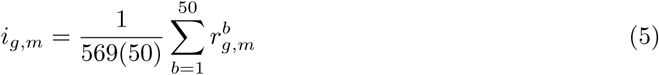

where the importance scores, *i*_*g,m*_, range from (0, 1], with 1 meaning that miRNA *m* is always the most important transcript for predicting mRNA *g*, and 0 meaning it is always the least important. Using this, we built a bipartite graph of importance where an edge between mRNA *g* and miRNA *m* indicates that its importance score *i*_*g,m*_ is among the top 2.5% of all importance scores. We repeated this for the top 5% and top 10% of importance scores too.

### 2.7 Gene set enrichment

Using the DE results which compared the paired primary and metastatic tumour samples, we ranked each mRNA gene based on the following formula:

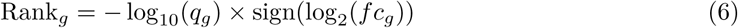

where *fc* is the fold-change and *q* the FDR-adjusted *p*-value as given by DESeq2. We then performed gene set enrichment analysis on this ranked gene list using GSEA (Version 3.0) [36] in PreRanked mode with classic enrichment and 1,000 permutations. We calculated enrichment scores for the **MSigDB Hallmarks** gene sets (Version 6.1) [36, 22, 23]. Note that the ranks used here differ from the non-parametric ranks used in “Bipartite graph analysis”.

## 3. Results and Discussion

### 3.1 mRNA as a cross-cancer metastasis biomarker

Differential expression (DE) analysis reveals 267 mRNA genes with a significant difference in abundance between the primary and metastatic pairs. Although these samples derive from several cancers, the significantly DE genes have a consistent direction of change across all of the involved cancers. Figure 1 shows the distribution of the relative log-fold change (LFC) of the metastatic tumour expression over its paired primary tumour expression. Owing to the paired design, we can infer that these differences likely arise from the evolution of cancer metastasis.

**Figure 1:**
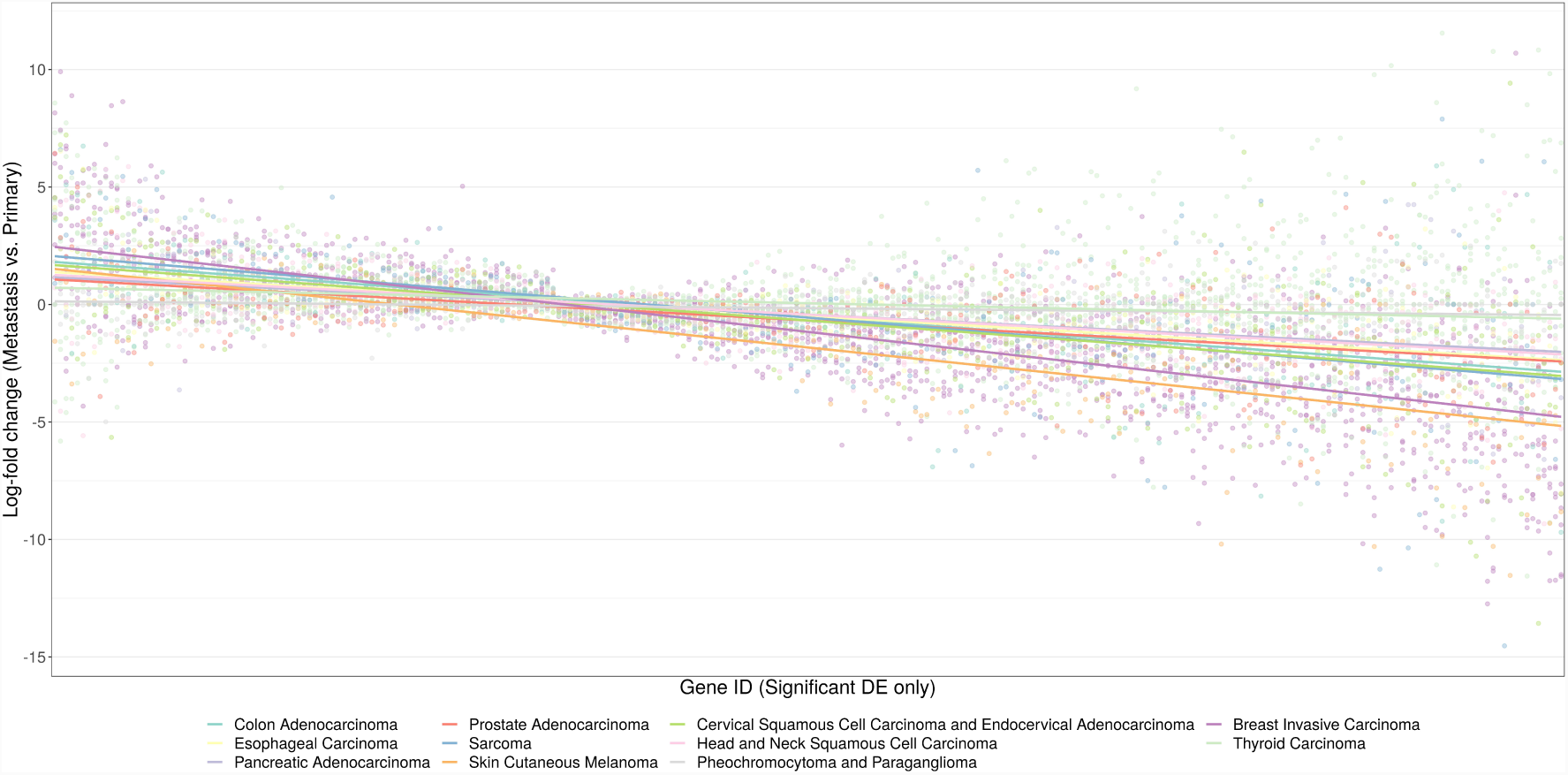
This figure shows the per-patient log-fold change (LFC) of metastatic expression over primary tumour expression (y-axis) for the significant differentially expressed (DE) gene (x-axis), coloured by cancer type. Here, we see that significant DE genes show a consistent expression signature across the 11 cancer types. Although some cancer types tend to have higher LFCs than others (e.g., Skin Cutaneous Melanoma), no cancer type deviates extremely from the general pattern, though individual outliers exist. Indeed, even for the cancer types with the lowest LFCs, the average LFC direction follows the same average trend as the cancer types with the highest LFCs. As such, these genes comprise a cross-cancer metastasis signature.

DE analysis identifies genes that associate with the outcome beyond chance. However, it does not tell us whether the genes would serve as useful biomarkers to differentiate metastatic and primary samples diagnostically. For this, we tested three classification algorithms with bootstrapped cross-validation to measure how accurately mRNA biomarkers can classify metastasis. Figure 2 shows the performance of the LASSO, random forest (RF), and support vector machine (SVM) classifiers. With an average 64.1%, 71.3%, and 70.1% accuracy for the top 64 genes, we can conclude that it is possible to accurately classify whether a cross-cancer sample is metastatic or primary based only on its gene expression signature.

**Figure 2:**
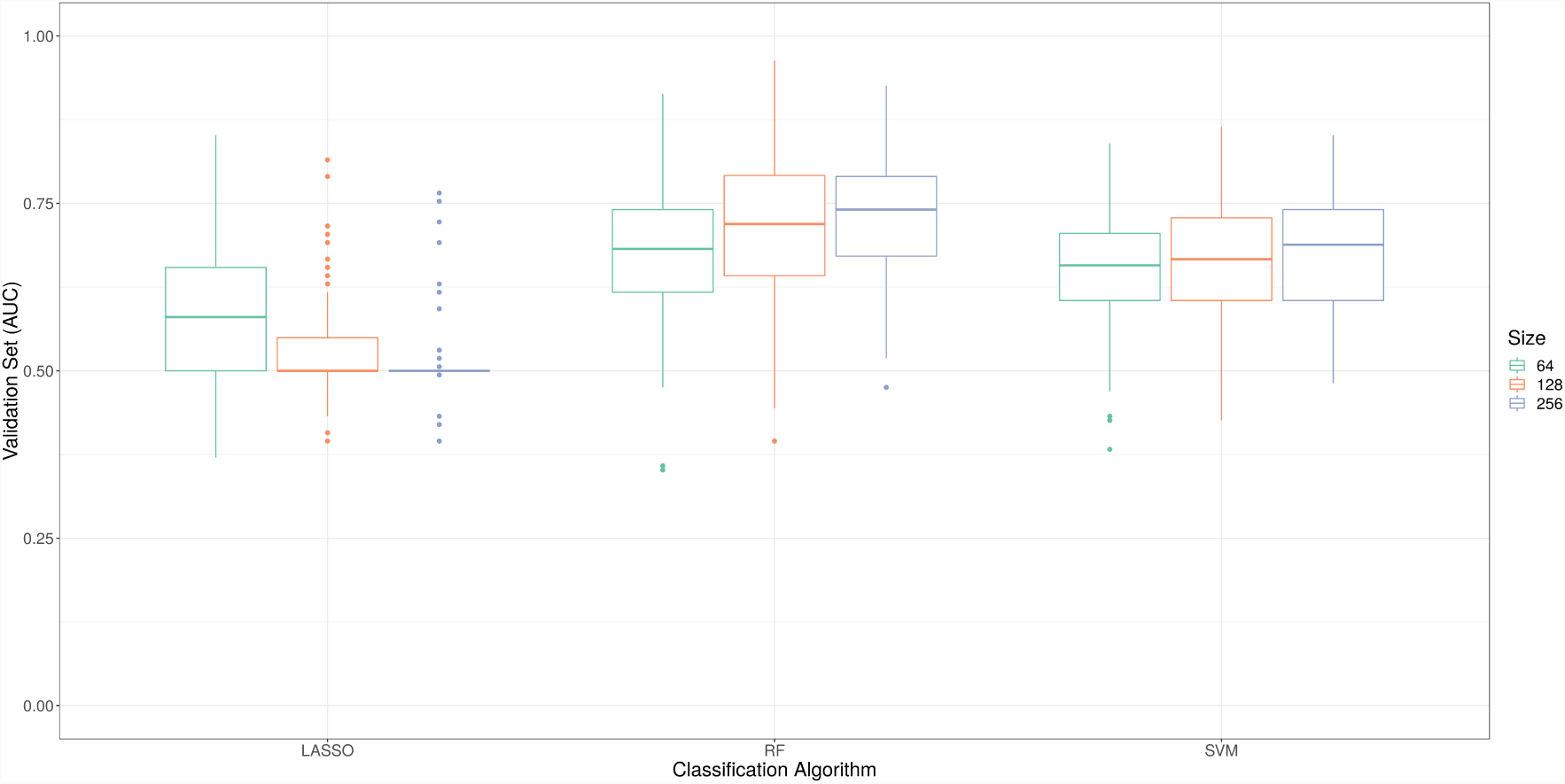
This figure shows the classification accuracy of metastatic status for paired tumour samples across 100 Monte-Carlo re-samplings (y-axis) for each classification model (x-axis) using only the mRNA expression. Boxplot colour refers to the top {*N* = 64, 128, 256} features as selected by Student’s t-test. Although the classifier size somewhat impacts performance, LASSO is always outperformed by random forest (RF) and support vector machine (SVM) regardless of classifier size. Empirically, RF appears to perform the best.

Using the DE results, we can also test whether higher-level pathway enrichments distinguish metastatic from primary tumours. From a gene set enrichment analysis (GSEA) of the **MSigDB Hallmarks** gene sets, we found 33 pathways that were significantly enriched or depleted (FDR-adjusted *p*-value < 0.05). Figure 3 shows the enrichment scores for these pathways, a number of which have a well-known involvement in metastatic processes, including *epithelial mesenchymal transition*, *G2M checkpoint*, *angiogenesis*, and several proliferative signalling pathways. Taken together, our results show that significant differences exist between the mRNA expression profile of primary and metastatic tumours, and that these differences are conserved across multiple cancer types. These differences not only have predictive value, but also reflect cellular processes that are known to drive metastatic progression.

**Figure 3:**
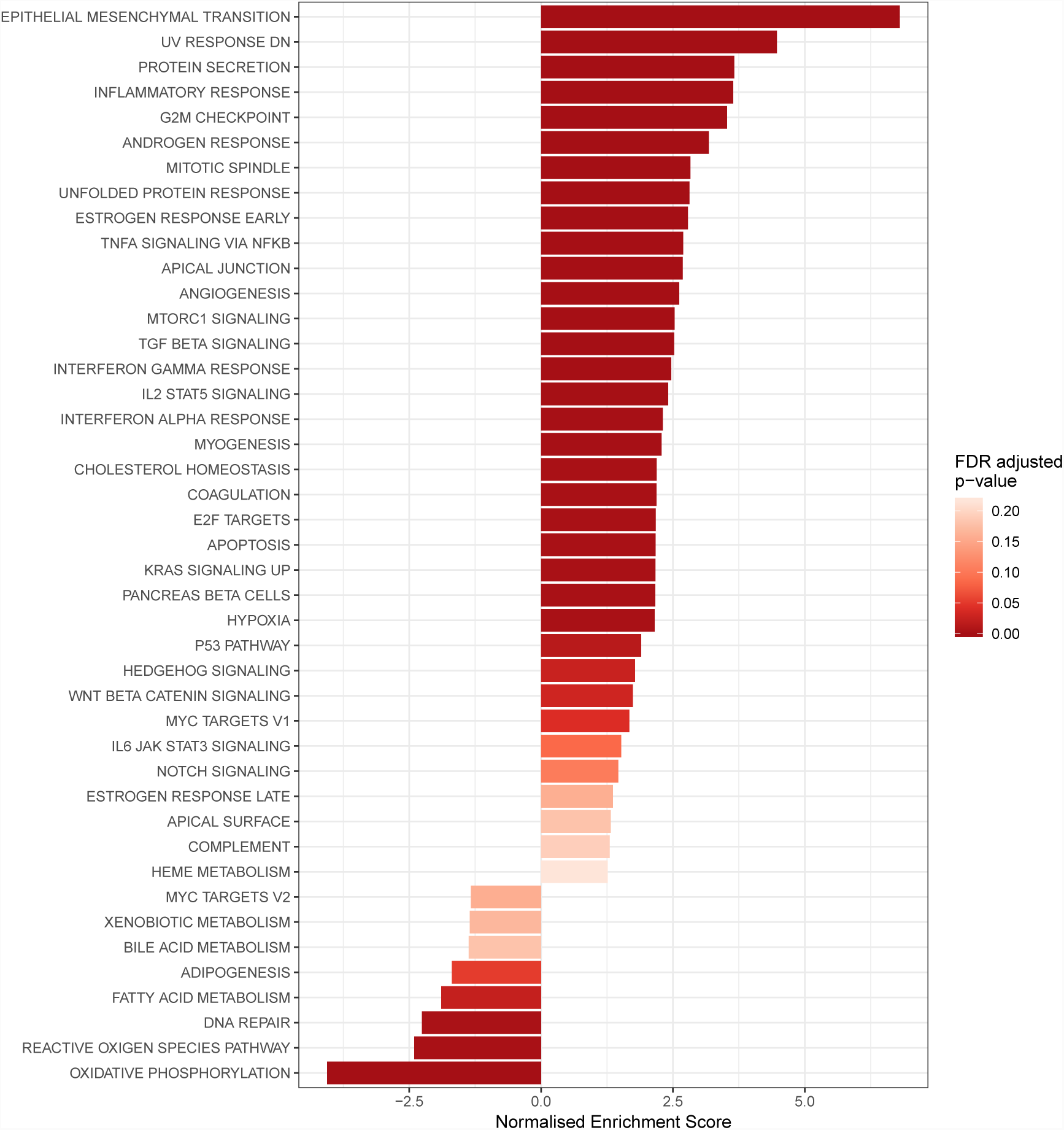
This figure shows the normalised enrichment scores from a gene set enrichment analysis (GSEA) of the mRNA differential expression (DE) results which compare metastatic samples with paired primary tumours. Using the **MSigDB Hallmarks** gene set, we find 29 enriched and 4 depleted pathway (FDR-adjusted *p*-value < 0.05). Each bar represents a pathway that is coloured by statistical significance, with a darker shade of red corresponding to a smaller *p*-value. Many of the pathways enriched here are involved in canonical cancer pathways.

### 3.2 microRNA as a cross-cancer metastasis biomarker

DE analysis uncovered no miRNA transcripts with a significant difference in abundance between the primary and metastatic pairs. We found this surprising considering the literature strongly suggests that miRNAs have an important role in cancer metastasis. Given our small sample size of 27 pairs, it is possible that this analysis was under-powered to detect DE miRNA transcripts. However, we did find several hundred significantly DE mRNA genes despite the larger FDR penalty applied to those results. One explanation for this finding is that miRNAs could have roles specific to the type of cancer undergoing metastasis, thus precluding the discovery of a cross-cancer signature. Alternatively, miRNAs may have more variance in general, necessitating larger sample sizes.

Although we found no significantly DE miRNA transcripts, it is still possible that a multivariate contribution of miRNA signatures could meaningfully differentiate metastatic from primary tumours. If so, we could interpret the predictive contribution made by the modelled miRNA transcripts directly. As above, we tested three classification algorithms with bootstrapped crossvalidation to measure how accurately miRNA biomarkers can classify metastasis. Figure 4 shows the performance of the LASSO, RF, and SVM classifiers. Despite the absence of significantly DE miRNA transcripts, we observed appreciable accuracy with the RF and SVM models. With an average 64.9% and 64.1% accuracy for the top 64 genes, we can conclude that it is possible to classify with moderate accuracy whether a cross-cancer sample is metastatic or primary based only on its miRNA expression signature. However, the miRNA signature does not discriminate the groups as well as the mRNA gene expression signature. These results agree with the existing literature which suggests miRNA signatures can be used to classify cancer, including malignancy [18].

**Figure 4:**
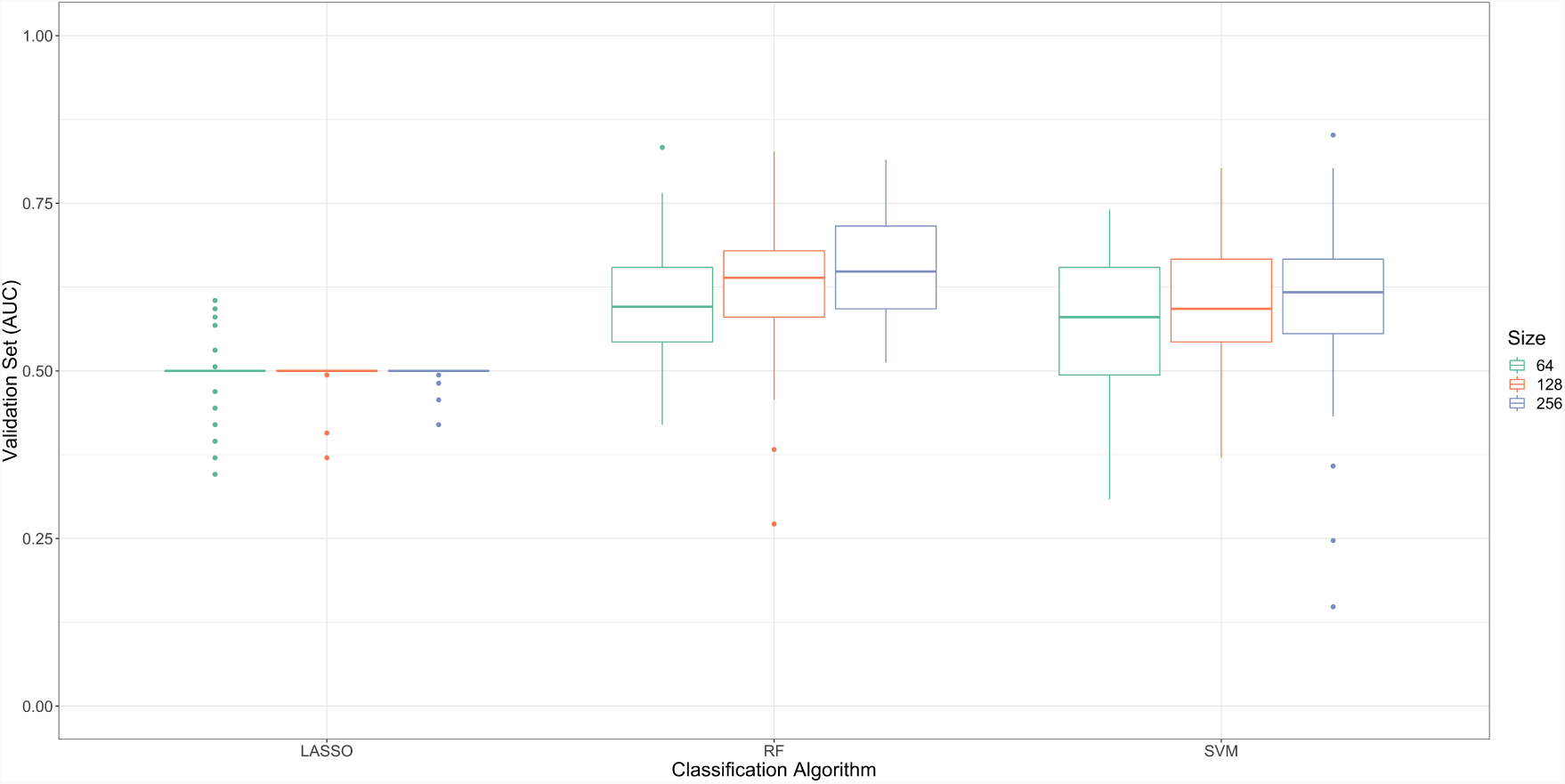
This figure shows the classification accuracy of metastatic status for paired tumour samples across 100 Monte-Carlo re-samplings (y-axis) for each classification model (x-axis) using only the mRNA expression. Boxplot colour refers to the top {*N* = 64, 128, 256} features as selected by Student’s t-test. Similar to Figure 2, classifier size somewhat impacts performance. However, LASSO fails to classify tumour status. Although random forest (RF) and support vector machine (SVM) do classify metastatic status based on miRNA expression, using mRNA expression works better.

Given that both mRNA and miRNA expression can accurately classify metastasis, we also tested whether the combination of these two feature sets further improves performance. However, we did not see any improvement over using mRNA expression alone (see Supplementary Information). This finding is consistent with the existing literature [41].

### 3.3 microRNA as a predictor of mRNA expression

Based on our understanding of how miRNA transcripts can act directly to regulate mRNA abundance, it seems plausible that we could accurately model mRNA expression in terms of miRNA expression. As such, we designed another cross-validation scheme to measure how accurately we can predict gene LFC based only on miRNA LFC, assessing significance through a permutation-based procedure. For this, we use a RF model because it tends to perform well for high-dimensional regression tasks while also allowing the analyst to interpret relative feature importance through “node purity”.

Figure 5 shows a bipartite graph that highlights the most important connections between miRNA and mRNA expression (with importance defined in “Bipartite graph analysis”). For this graph, the presence of a mRNA gene node indicates that its metastatic dysregulation can be predicted by miRNA dysregulation (FDR-adjusted *p <* 0.05). Twenty-eight genes satisfy this criteria, as presented in Table 1. We interpret this to mean that the expression of these mRNA are somehow linked to, and possibly influenced by, the expression of the associated miRNAs.

**Table 1:** This table shows the 28 mRNAs whose metastatic log-fold change (LFC) is predicted by the metastatic LFC of miRNAs significantly better than chance. In this table, we present the HGNC symbol for the Ensebl gene ID, the permuted root mean squared error (RMSE) of the regression, the observed RMSE of the regression, and the percent difference between the permuted and observed RMSEs. We also report *p*-values for a one-tailed Wilcoxon Rank-Sum test comparing the permuted and observed RMSEs. To control the false discovery rate, we adjusted the *p*-values using the Benjamini-Hochberg procedure.

| Ensembl ID | Gene Symbol | Permuted RMSE | Observed RMSE | Percent Difference | $p$-value | BH $p$-value |
| --- | --- | --- | --- | --- | --- | --- |
| ENSG00000110427 | KIAA1549L | 1.6989 | 1.4224 | 19.4355 | 0.0000 | 0.0000 |
| ENSG00000142661 | MYOM3 | 1.4599 | 1.2425 | 17.4914 | 0.0000 | 0.0000 |
| ENSG00000167483 | FAM129C | 1.9533 | 1.6211 | 20.4927 | 0.0000 | 0.0000 |
| ENSG00000168071 | CCDC88B | 0.8564 | 0.6996 | 22.4078 | 0.0000 | 0.0000 |
| ENSG00000232679 | LINC01705 | 1.6862 | 1.4530 | 16.0480 | 0.0000 | 0.0000 |
| ENSG00000134363 | FST | 1.4929 | 1.2823 | 16.4227 | 0.0000 | 0.0001 |
| ENSG00000137809 | ITGA11 | 1.2159 | 1.0114 | 20.2262 | 0.0000 | 0.0001 |
| ENSG00000165617 | DACT1 | 0.8431 | 0.7272 | 15.9438 | 0.0000 | 0.0001 |
| ENSG00000176971 | FIBIN | 1.8796 | 1.4608 | 28.6640 | 0.0000 | 0.0002 |
| ENSG00000189320 | FAM180A | 1.8069 | 1.4439 | 25.1400 | 0.0000 | 0.0002 |
| ENSG00000164107 | HAND2 | 1.9638 | 1.7251 | 13.8377 | 0.0000 | 0.0003 |
| ENSG00000156103 | MMP16 | 1.3396 | 1.1962 | 11.9896 | 0.0000 | 0.0006 |
| ENSG00000122986 | HVCN1 | 0.8464 | 0.7609 | 11.2370 | 0.0000 | 0.0009 |
| ENSG00000145824 | CXCL14 | 3.6598 | 3.1571 | 15.9248 | 0.0000 | 0.0009 |
| ENSG00000157551 | KCNJ15 | 1.1121 | 0.9751 | 14.0453 | 0.0000 | 0.0013 |
| ENSG00000204262 | COL5A2 | 1.2158 | 1.0178 | 19.4585 | 0.0000 | 0.0020 |
| ENSG00000075035 | WSCD2 | 1.6657 | 1.4472 | 15.0967 | 0.0000 | 0.0021 |
| ENSG00000180448 | ARHGAP45 | 0.6604 | 0.5820 | 13.4832 | 0.0000 | 0.0026 |
| ENSG00000166923 | GREM1 | 1.7331 | 1.4957 | 15.8748 | 0.0000 | 0.0031 |
| ENSG00000186081 | KRT5 | 3.6900 | 3.1928 | 15.5709 | 0.0000 | 0.0042 |
| ENSG00000171812 | COL8A2 | 1.1490 | 1.0070 | 14.1091 | 0.0000 | 0.0068 |
| ENSG00000145423 | SFRP2 | 3.2499 | 2.6971 | 20.4931 | 0.0000 | 0.0080 |
| ENSG00000133110 | POSTN | 1.7425 | 1.5359 | 13.4543 | 0.0000 | 0.0105 |
| ENSG00000174099 | MSRB3 | 0.7309 | 0.6276 | 16.4519 | 0.0000 | 0.0111 |
| ENSG00000227039 | ITGB2-AS1 | 0.9453 | 0.8424 | 12.2113 | 0.0001 | 0.0162 |
| ENSG00000105989 | WNT2 | 2.5691 | 2.2515 | 14.1059 | 0.0001 | 0.0167 |
| ENSG00000180044 | C3orf80 | 1.2539 | 1.0542 | 18.9402 | 0.0001 | 0.0224 |
| ENSG00000145244 | CORIN | 1.1041 | 0.9839 | 12.2145 | 0.0002 | 0.0470 |

**Figure 5:**
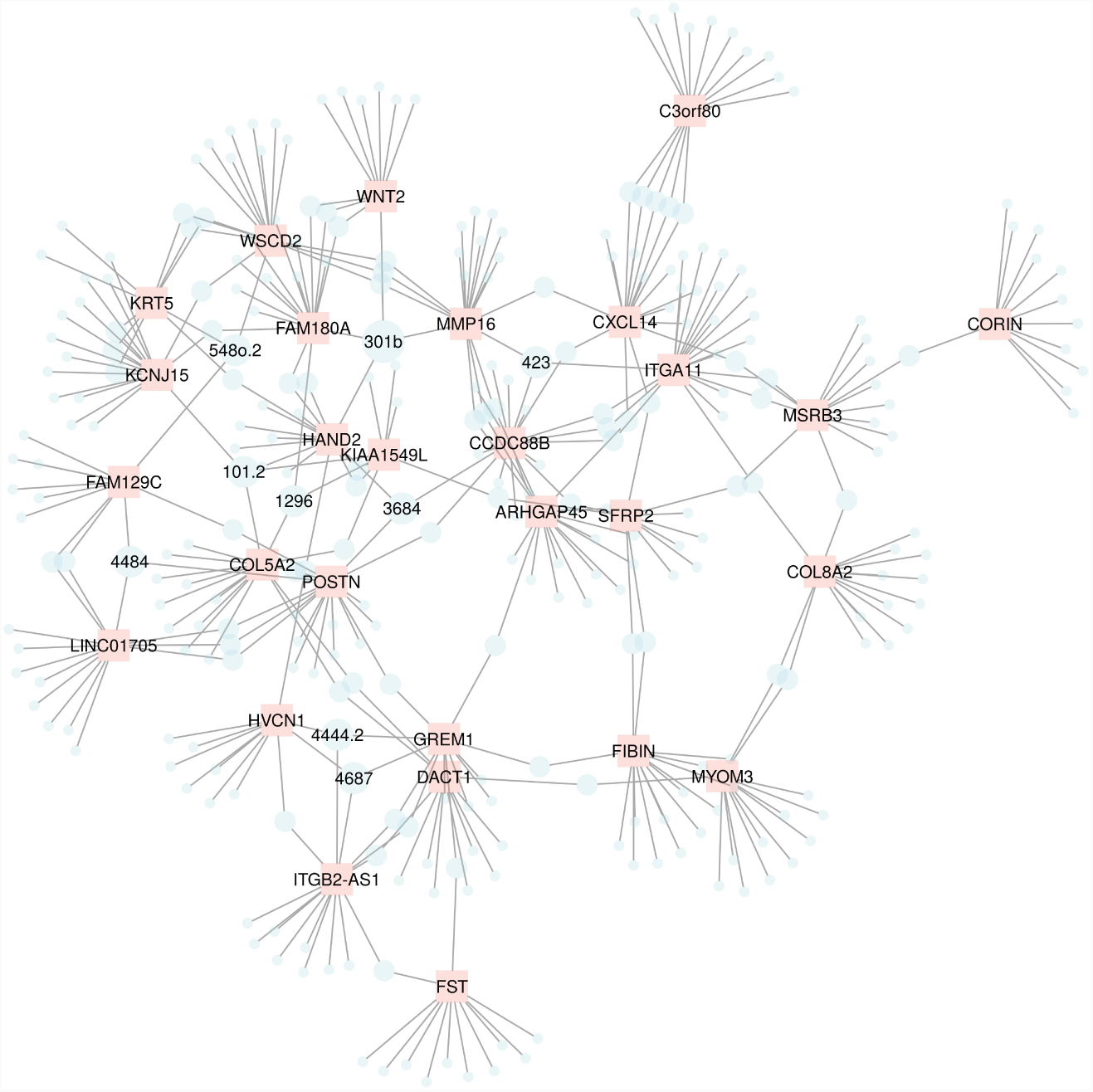
This figure shows the top associations (edges), as ranked by node purity, between miRNAs (blue nodes) and mRNAs (pink nodes). Included in this figure are 28 mRNAs whose metastatic log-fold change (LFC) can be predicted by the metastatic LFC of miRNA (FDR-adjusted *p <* 0.05). The labelled miRNA transcripts are important in predicting the dysregulation of at least 3 mRNAs. Some of these labelled miRNAs have been previously implicated in cancer metastasis or progression, including miR-301b, miR-1296, and miR-423. The complete edge list is provided in the Supplementary Information. Note that for clarify of visualization, we removed the “miR-” prefix from the miRNA node labels. Interestingly, most of these genes are not characterised within the **MSigDB Hallmarks** or **KEGG** gene sets.

Edges, on the other hand, indicate that a change in the metastatic expression of that miRNA helps predict the metastatic dysregulation of that mRNA gene. Here, we see 9 miRNAs that have an important contribution in predicting the differential expression of at least 3 mRNAs. Of these, miR-301b, a member of the part of the miR-130 family [11], associates with the metastatic signature of 4 mRNAs. This miRNA has been found up-regulated in malignant prostate cancer samples [11], and is associated with worse breast cancer outcomes [7]. Indeed, miR-301b is considered part of a super-family of pan-cancer oncomiRs, along with the miR-17, -19, -130, -210 -18 and -455 families, some of which also appear as lesser nodes in Figure 5 (i.e., -17, -19, -130, and -18) [16]. Meanwhile, two miRNAs within the miR-130 family, miR-301a and miR-130a, associate with 1 and 2 genes, respectively. Although the choice to analyse the top 2.5% of important edges was arbitrary, the combined degree of the miR-130 family is still larger than that of any single miR when looking at the top 5% and top 10% of edges (with 11 and 16 associations respectively). Taken together, our data suggest that a LFC of miR-130 transcripts in a metastatic sample is associated with a LFC of select mRNAs in the same metastatic sample.

Among the most “important” miRNAs, we also find miR-1296 to be associated with the metastatic signature of 3 mRNAs. As above, miR-1296 is associated with cancer and metastasis, with one study finding it down-regulated in gastric cancer (compared with adjacent normal tissue) and even more down-regulated in lymph node metastases (compared with gastric cancer) [34]. Meanwhile, induction of miR-1296 was found to induce apoptosis in triple negative breast cancer [28]. Although the other miRNAs presented in Figure 5 have not yet been associated with metastasis *per se*, several have been linked to cancer more generally, including miR-423 which is found to promote tumour progression in laryngeal cancer [15], and miR-101-2 and miR-4687 which are found to be associated with gastric cancer [31] and breast cancer [25] respectively. Confirmatory studies are needed to determine whether these miRNAs promote oncogenesis through the regulation of genes predicted by our random forest model.

## 4. Summary

The miRNA-mRNA regulatory axis remains an important facet of cancer research with potential diagnostic and therapeutic implications. For this reason, we used paired primary and metastatic samples to explore the miRNA-mRNA regulatory axis across 11 cancers from three perspectives. First, we identified a cross-cancer mRNA signal capable of differentiating primary tumours from metastatic. This signal contains biomarkers enriched for canonical cancer pathways including *epithelial mesenchymal transition* and *G2M checkpoint*. Second, we also identified a cross-cancer miRNA signal capable of differentiating primary and metastatic samples, although this signal was not as strong as the mRNA signal. Finally, we integrated the mRNA and miRNA data to model how well the metastasis-associated dysregulation of miRNA can predict the metastasis-associated dysregulation of mRNA. From this analysis, we identified several miRNAs whose dysregulation alone can predict mRNA dysregulation, including miR-301b, miR-1296, and miR-423, previously linked to cancer metastasis or progression. These discoveries all survive rigorous statistical correction despite small sample sizes, highlighting the value of paired study designs. Nevertheless, experimental validation is needed to determine whether the discovered miRNA-mRNA associations are causal.

## 5. Declarations

### 5.1 Ethics approval and consent to participate

Not applicable.

### 5.2 Consent for publication

Not applicable.

### 5.3 Availability of data and material

All data and scripts are publicly available.

### 5.4 Competing interests

No authors have competing interests.

### 5.5 Funding

Not applicable.

### 5.6 Authors’ contributions

SCL and TPQ reviewed the literature, designed the project, performed the analyses, and drafted the manuscript. AQ reviewed the literature and helped draft the manuscript. All authors edited and approved the final manuscript.

## Supporting information

## 5.7 Acknowledgements

Not applicable.

## References

[1] U. Alon, N. Barkai, D. A. Notterman, K. Gish, S. Ybarra, D. Mack, and A. J. Levine. Broad patterns of gene expression revealed by clustering analysis of tumor and normal colon tissues probed by oligonucleotide arrays. Proceedings of the National Academy of Sciences of the United States of America, 96(12):6745–6750, June 1999.

[2] Simon Anders and Wolfgang Huber. Differential expression analysis for sequence count data. Genome Biology, 11:R106, October 2010.

[3] Virginie Armand-Labit and Anne Pradines. Circulating cell-free microRNAs as clinical cancer biomarkers. Biomolecular Concepts, 8(2):61–81, May 2017.

[4] Yoav Benjamini and Yosef Hochberg. Controlling the False Discovery Rate: A Practical and Powerful Approach to Multiple Testing. Journal of the Royal Statistical Society. Series B (Methodological), 57(1):289–300, 1995.

[5] C. I. Bliss and R. A. Fisher. Fitting the Negative Binomial Distribution to Biological Data. Biometrics, 9(2):176–200, 1953.

[6] Robert R. J. Coebergh van den Braak, Anieta M. Sieuwerts, Zarina S. Lalmahomed, Marcel Smid, Saskia M. Wilting, Sandra I. Bril, Shanshan Xiang, Michelle van der Vlugt-Daane, Vanja de Weerd, Anne van Galen, Katharina Biermann, J. Han J. M. van Krieken, Wigard P. Kloosterman, John A. Foekens, John W. M. Martens, and Jan N. M. IJzermans. Confirmation of a metastasis-specific microRNA signature in primary colon cancer. Scientific Reports, 8(1):5242, March 2018.

[7] D. Cheng, H. He, and B. Liang. A three-microRNA signature predicts clinical outcome in breast cancer patients. European Review for Medical and Pharmacological Sciences, 22(19):6386–6395, October 2018.

[8] Ajay Francis Christopher, Raman Preet Kaur, Gunpreet Kaur, Amandeep Kaur, Vikas Gupta, and Parveen Bansal. MicroRNA therapeutics: Discovering novel targets and developing specific therapy. Perspectives in Clinical Research, 7(2):68–74, 2016.

[9] Andy Chu, Gordon Robertson, Denise Brooks, Andrew J. Mungall, Inanc Birol, Robin Coope, Yussanne Ma, Steven Jones, and Marco A. Marra. Large-scale profiling of microRNAs for The Cancer Genome Atlas. Nucleic Acids Research, 44(1):e3–e3, January 2016.

[10] Antonio Colaprico, Claudia Cava, Gloria Bertoli, Gianluca Bontempi, and Isabella Castiglioni. Integrative Analysis with Monte Carlo Cross-Validation Reveals miRNAs Regulating Pathways Cross-Talk in Aggressive Breast Cancer. BioMed research international, 2015:831314, 2015.

[11] Rafael Sebastián Fort, Cecilia Mathó, Carolina Oliveira-Rizzo, Beatriz Garat, José Roberto Sotelo-Silveira, and María Ana Duhagon. An integrated view of the role of miR-130b/301b miRNA cluster in prostate cancer. Experimental Hematology & Oncology, 7:10, 2018.

[12] Jerome Friedman, Trevor Hastie, and Rob Tibshirani. Regularization Paths for Generalized Linear Models via Coordinate Descent. Journal of Statistical Software, 33(1):1–22, 2010.

[13] Michela Garofalo and Carlo M. Croce. microRNAs: Master regulators as potential therapeutics in cancer. Annual Review of Pharmacology and Toxicology, 51:25–43, 2011.

[14] T. R. Golub, D. K. Slonim, P. Tamayo, C. Huard, M. Gaasenbeek, J. P. Mesirov, H. Coller, M. L. Loh, J. R. Downing, M. A. Caligiuri, C. D. Bloomfield, and E. S. Lander. Molecular classification of cancer: class discovery and class prediction by gene expression monitoring. Science (New York, N.Y.), 286(5439):531–537, October 1999.

[15] Guofang Guan, Dejun Zhang, Ying Zheng, Lianji Wen, Duojiao Yu, Yanqing Lu, and Yan Zhao. microRNA-423-3p promotes tumor progression via modulation of AdipoR2 in laryngeal carcinoma. International Journal of Clinical and Experimental Pathology, 7(9):5683–5691, August 2014.

[16] Mark P. Hamilton, Kimal Rajapakshe, Sean M. Hartig, Boris Reva, Michael D. McLellan, Cyriac Kandoth, Li Ding, Travis I. Zack, Preethi H. Gunaratne, David A. Wheeler, Cristian Coarfa, and Sean E. McGuire. Identification of a pan-cancer oncogenic microRNA superfamily anchored by a central core seed motif. Nature Communications, 4:2730, November 2013.

[17] Steven R. Head, H. Kiyomi Komori, Sarah A. LaMere, Thomas Whisenant, Filip Van Nieuwerburgh, Daniel R. Salomon, and Phillip Ordoukhanian. Library construction for next-generation sequencing: Overviews and challenges. BioTechniques, 56(2):61–passim, February 2014.

[18] Marilena V. Iorio and Carlo M. Croce. MicroRNA dysregulation in cancer: diagnostics, monitoring and therapeutics. A comprehensive review. EMBO Molecular Medicine, 4(3):143–159, March 2012.

[19] Limin Li, Jing Zhang, Wenli Diao, Dong Wang, Yao Wei, Chen-Yu Zhang, and Ke Zen. MicroRNA-155 and MicroRNA-21 promote the expansion of functional myeloid-derived suppressor cells. Journal of Immunology (Baltimore, Md.: 1950), 192(3):1034–1043, February 2014.

[20] Wenhua Li, Jinjia Chang, Duo Tong, Junjie Peng, Dan Huang, Weijian Guo, Wen Zhang, and Jin Li. Differential microRNA expression profiling in primary tumors and matched liver metastasis of patients with colorectal cancer. Oncotarget, 8(22):35783–35791, May 2017.

[21] Andy Liaw and Matthew Wiener. Classification and Regression by randomForest. R News, 2(3):18–22, 2002.

[22] A. Liberzon, A. Subramanian, R. Pinchback, H. Thorvaldsdottir, P. Tamayo, and J. P. Mesirov. Molecular signatures database (MSigDB) 3.0. Bioinformatics, 27(12):1739–1740, 6 2011.

[23] Arthur Liberzon, Chet Birger, Helga Thorvaldsdóttir, Mahmoud Ghandi, Jill P. Mesirov, and Pablo Tamayo. The Molecular Signatures Database Hallmark Gene Set Collection. Cell Systems, 1(6):417–425, 12 2015.

[24] Weiyang Lou, Jingxing Liu, Yanjia Gao, Guansheng Zhong, Danni Chen, Jiaying Shen, Chang Bao, Liang Xu, Jie Pan, Junchi Cheng, Bisha Ding, and Weimin Fan. MicroRNAs in cancer metastasis and angiogenesis. Oncotarget, 8(70):115787–115802, December 2017.

[25] Diana V. Maltseva, Vladimir V. Galatenko, Timur R. Samatov, Svetlana O. Zhikrivetskaya, Nadezhda A. Khaustova, Ilya N. Nechaev, Maxim U. Shkurnikov, Alexey E. Lebedev, Irina A. Mityakina, Andrey D. Kaprin, Udo Schumacher, and Alexander G. Tonevitsky. miRNome of inflammatory breast cancer. BMC Research Notes, 7(1):871, December 2014.

[26] David Meyer, Evgenia Dimitriadou, Kurt Hornik, Andreas Weingessel, and Friedrich Leisch. e1071: Misc Functions of the Department of Statistics, Probability Theory Group (Formerly: E1071), TU Wien. 2017.

[27] Keith Noto, Saeed Majidi, Andrea G. Edlow, Heather C. Wick, Diana W. Bianchi, and Donna K. Slonim. CSAX: Characterizing Systematic Anomalies in eXpression Data. Journal of Computational Biology, 22(5):402–413, May 2015.

[28] Binh Phan, Shahana Majid, Sarah Ursu, David de Semir, Mehdi Nosrati, Vladimir Bezrookove, Mohammed Kashani-Sabet, and Altaf A. Dar. Tumor suppressor role of microRNA-1296 in triple-negative breast cancer. Oncotarget, 7(15):19519–19530, January 2016.

[29] Thomas Quinn, Daniel Tylee, and Stephen Glatt. exprso: an R-package for the rapid implementation of machine learning algorithms. F1000Research, 5:2588, December 2017.

[30] Thomas P. Quinn, Thin Nguyen, Samuel C. Lee, and Svetha Venkatesh. Cancer as a tissue anomaly: classifying tumor transcriptomes based only on healthy data. bioRxiv, page 426395, September 2018.

[31] Ismael Riquelme, Oscar Tapia, Pamela Leal, Alejandra Sandoval, Matthew G. Varga, Pablo Letelier, Kurt Buchegger, Carolina Bizama, Jaime A. Espinoza, Richard M. Peek, Juan Carlos Araya, and Juan Carlos Roa. miR-101-2, miR-125b-2 and miR-451a act as potential tumor suppressors in gastric cancer through regulation of the PI3k/AKT/mTOR pathway. Cellular Oncology, 39(1):23–33, February 2016.

[32] Mark D. Robinson, Davis J. McCarthy, and Gordon K. Smyth. edgeR: a Bioconductor package for differential expression analysis of digital gene expression data. Bioinformatics, 26(1):139–140, January 2010.

[33] Zachary Schrank, Nabiha Khan, Chike Osude, Sanjana Singh, Rachel J. Miller, Collin Merrick, Alexander Mabel, Adijan Kuckovic, and Neelu Puri. Oligonucleotides Targeting Telomeres and Telomerase in Cancer. Molecules (Basel, Switzerland), 23(9), September 2018.

[34] Xia Shan, Wei Wen, Danxia Zhu, Ting Yan, Wenfang Cheng, Zebo Huang, Lan Zhang, Huo Zhang, Tongshan Wang, Wei Zhu, Yichao Zhu, and Jun Zhu. miR 1296-5p Inhibits the Migration and Invasion of Gastric Cancer Cells by Repressing ERBB2 Expression. PLOS ONE, 12(1):e0170298, January 2017.

[35] Chikako Shibata, Motoyuki Otsuka, Takahiro Kishikawa, Takeshi Yoshikawa, Motoko Ohno, Akemi Takata, and Kazuhiko Koike. Current status of miRNA-targeting therapeutics and preclinical studies against gastroenterological carcinoma. Molecular and cellular therapies, 1, December 2013.

[36] Aravind Subramanian, Pablo Tamayo, Vamsi K. Mootha, Sayan Mukherjee, Benjamin L. Ebert, Michael A. Gillette, Amanda Paulovich, Scott L. Pomeroy, Todd R. Golub, Eric S. Lander, and Jill P. Mesirov. Gene set enrichment analysis: A knowledge-based approach for interpreting genome-wide expression profiles. Proceedings of the National Academy of Sciences, 102(43):15545–15550, October 2005.

[37] Laura J. van ’t Veer, Hongyue Dai, Marc J. van de Vijver, Yudong D. He, Augustinus A. M. Hart, Mao Mao, Hans L. Peterse, Karin van der Kooy, Matthew J. Marton, Anke T. Witteveen, George J. Schreiber, Ron M. Kerkhoven, Chris Roberts, Peter S. Linsley, René Bernards, and Stephen H. Friend. Gene expression profiling predicts clinical outcome of breast cancer. Nature, 415(6871):530–536, January 2002.

[38] John N. Weinstein, Eric A. Collisson, Gordon B. Mills, Kenna M. Shaw, Brad A. Ozenberger, Kyle Ellrott, Ilya Shmulevich, Chris Sander, and Joshua M. Stuart. The Cancer Genome Atlas Pan-Cancer Analysis Project. Nature genetics, 45(10):1113–1120, October 2013.

[39] Frank Wilcoxon. Individual Comparisons by Ranking Methods. Biometrics Bulletin, 1(6):80–83, 1945.

[40] Tingzhen Yan, Shiyong Zhu, Jing Zhang, Gongbiao Lu, Chaoliang Lv, Yanchun Wei, and Minghua Luo. MicroRNA-944 targets vascular endothelial growth factor to inhibit cell proliferation and invasion in osteosarcoma. Molecular Medicine Reports, October 2018.

[41] Qing Zhao, Xingjie Shi, Yang Xie, Jian Huang, BenChang Shia, and Shuangge Ma. Combining multidimensional genomic measurements for predicting cancer prognosis: observations from TCGA. Briefings in Bioinformatics, 16(2):291–303, March 2015.

