## Supplementary material for "A cross-cancer metastasis signature in the microRNA-mRNA axis of paired tissue samples"

**Abstract**

In the progression of cancer, cells acquire genetic mutations that cause uncontrolled growth. Over time, the primary tumour may undergo additional mutations that allow for the cancerous cells to spread throughout the body as metastases. Since metastatic development typically results in markedly worse patient outcomes, research into the identity and function of metastasis-associated biomarkers could eventually translate into clinical diagnostics or novel therapeutics. Although the general processes underpinning metastatic progression are understood, no consistent nor clear cross-cancer biomarker profile has yet emerged. However, the literature suggests that some microRNAs (miRNAs) may play an important role in the metastatic progression of several cancer types. Using a subset of The Cancer Genome Atlas (TCGA) data, we performed an integrated analysis of mRNA and miRNA expression with paired metastatic and primary tumour samples to interrogate how the miRNA-mRNA regulatory axis influences metastatic progression. From this, we successfully built mRNA- and miRNA-specific classifiers that can discriminate pairs of metastatic and primary samples across 11 cancer types. In addition, we identified a number of miRNAs whose metastasis-associated dysregulation could predict mRNA metastasis-associated dysregulation. Among the most predictive miRNAs, we found several previously implicated in cancer progression, including miR-301b, miR-1296, and miR-423. Taken together, our results suggest that cross-cancer metastatic samples have unique biomarker signatures when compared with paired primary tumours, and that these miRNA biomarkers can be used to predict both metastatic status and mRNA expression.

| Sample | Patient | Disease type | Tumour type | Primary diagnosis | Tumour stage | Age at diagnosis | Vital Status | Days to death | Gender | Year of birth | Race |
|---|---|---|---|---|---|---|---|---|---|---|---|
| TCGA-AC-A6IX-01A | TCGA-AC-A6IX | Breast Invasive Carcinoma | Primary solid Tumor | C50.9 | stage iiic | 17969 | alive | NA | female | 1964 | white |
| TCGA-AC-A6IX-06A | TCGA-AC-A6IX | Breast Invasive Carcinoma | Metastatic | C50.9 | stage iiic | 17969 | alive | NA | female | 1964 | white |
| TCGA-BH-A18V-01A | TCGA-BH-A18V | Breast Invasive Carcinoma | Primary solid Tumor | C50.4 | stage iib | 17682 | dead | 1556 | female | 1953 | white |
| TCGA-BH-A18V-06A | TCGA-BH-A18V | Breast Invasive Carcinoma | Metastatic | C50.4 | stage iib | 17682 | dead | 1556 | female | 1953 | white |
| TCGA-BH-A1ES-01A | TCGA-BH-A1ES | Breast Invasive Carcinoma | Primary solid Tumor | C50.9 | stage iib | 13138 | dead | 3462 | female | 1964 | white |
| TCGA-BH-A1ES-06A | TCGA-BH-A1ES | Breast Invasive Carcinoma | Metastatic | C50.9 | stage iib | 13138 | dead | 3462 | female | 1964 | white |
| TCGA-BH-A1FE-01A | TCGA-BH-A1FE | Breast Invasive Carcinoma | Primary solid Tumor | C50.9 | stage iib | 11650 | dead | 2273 | female | 1967 | white |
| TCGA-BH-A1FE-06A | TCGA-BH-A1FE | Breast Invasive Carcinoma | Metastatic | C50.9 | stage iib | 11650 | dead | 2273 | female | 1967 | white |
| TCGA-DE-A4MD-01A | TCGA-DE-A4MD | Thyroid Carcinoma | Primary solid Tumor | C73 | stage iva | 26179 | alive | NA | male | 1940 | white |
| TCGA-DE-A4MD-06A | TCGA-DE-A4MD | Thyroid Carcinoma | Metastatic | C73 | stage iva | 26179 | alive | NA | male | 1940 | white |
| TCGA-E2-A15A-01A | TCGA-E2-A15A | Breast Invasive Carcinoma | Primary solid Tumor | C50.9 | stage iiic | 16750 | alive | NA | female | 1964 | white |

| | | | | | | | | | | | |
|---|---|---|---|---|---|---|---|---|---|---|---|
| TCGA-E2-A15A-06A | TCGA-E2-A15A | Breast Invasive Carcinoma | Metastatic | C50.9 | stage iiic | 16750 | alive | NA | female | 1964 | white |
| TCGA-E2-A15E-01A | TCGA-E2-A15E | Breast Invasive Carcinoma | Primary solid Tumor | C50.9 | stage iia | 14894 | alive | NA | female | 1969 | white |
| TCGA-E2-A15E-06A | TCGA-E2-A15E | Breast Invasive Carcinoma | Metastatic | C50.9 | stage iia | 14894 | alive | NA | female | 1969 | white |
| TCGA-E2-A15K-01A | TCGA-E2-A15K | Breast Invasive Carcinoma | Primary solid Tumor | C50.9 | stage iib | 21514 | alive | NA | female | 1952 | white |
| TCGA-E2-A15K-06A | TCGA-E2-A15K | Breast Invasive Carcinoma | Metastatic | C50.9 | stage iib | 21514 | alive | NA | female | 1952 | white |
| TCGA-EM-A2CS-01A | TCGA-EM-A2CS | Thyroid Carcinoma | Primary solid Tumor | C73 | stage iva | 18990 | alive | NA | female | 1960 | not reported |
| TCGA-EM-A2CS-06A | TCGA-EM-A2CS | Thyroid Carcinoma | Metastatic | C73 | stage iva | 18990 | alive | NA | female | 1960 | not reported |
| TCGA-EM-A2P1-01A | TCGA-EM-A2P1 | Thyroid Carcinoma | Primary solid Tumor | C73 | stage i | 12320 | alive | NA | male | 1978 | not reported |
| TCGA-EM-A2P1-06A | TCGA-EM-A2P1 | Thyroid Carcinoma | Metastatic | C73 | stage i | 12320 | alive | NA | male | 1978 | not reported |
| TCGA-EM-A3FQ-01A | TCGA-EM-A3FQ | Thyroid Carcinoma | Primary solid Tumor | C73 | stage i | 7176 | alive | NA | female | 1992 | not reported |
| TCGA-EM-A3FQ-06A | TCGA-EM-A3FQ | Thyroid Carcinoma | Metastatic | C73 | stage i | 7176 | alive | NA | female | 1992 | not reported |
| TCGA-EM-A3SU-01A | TCGA-EM-A3SU | Thyroid Carcinoma | Primary solid Tumor | C73 | stage i | 15768 | alive | NA | female | 1968 | not reported |
| TCGA-EM-A3SU-06A | TCGA-EM-A3SU | Thyroid Carcinoma | Metastatic | C73 | stage i | 15768 | alive | NA | female | 1968 | not reported |
| TCGA-ER-A2NF-01A | TCGA-ER-A2NF | Skin Cutaneous Melanoma | Primary solid Tumor | C44.4 | stage iiib | 19499 | dead | 877 | male | 1957 | white |

| | | | | | | | | | | | |
|---|---|---|---|---|---|---|---|---|---|---|---|
| TCGA-ER-A2NF-06A | TCGA-ER-A2NF | Skin Cutaneous Melanoma | Metastatic | C44.4 | stage iiib | 19499 | dead | 877 | male | 1957 | white |
| TCGA-HM-A6W2-01A | TCGA-HM-A6W2 | Cervical Squamous Cell Carcinoma and Endocervical Adenocarcinoma | Primary solid Tumor | C56.9 | not reported | 12775 | alive | NA | female | 1979 | black or african american |
| TCGA-HM-A6W2-06A | TCGA-HM-A6W2 | Cervical Squamous Cell Carcinoma and Endocervical Adenocarcinoma | Metastatic | C56.9 | not reported | 12775 | alive | NA | female | 1979 | black or african american |
| TCGA-HZ-A9TJ-01A | TCGA-HZ-A9TJ | Pancreatic Adenocarcinoma | Primary solid Tumor | C25.2 | stage iv | 25849 | alive | NA | male | 1943 | white |
| TCGA-HZ-A9TJ-06A | TCGA-HZ-A9TJ | Pancreatic Adenocarcinoma | Metastatic | C25.2 | stage iv | 25849 | alive | NA | male | 1943 | white |
| TCGA-J8-A3O2-01A | TCGA-J8-A3O2 | Thyroid Carcinoma | Primary solid Tumor | C73 | stage i | 14339 | alive | NA | male | 1972 | white |
| TCGA-J8-A3O2-06A | TCGA-J8-A3O2 | Thyroid Carcinoma | Metastatic | C73 | stage i | 14339 | alive | NA | male | 1972 | white |
| TCGA-J8-A3YH-01A | TCGA-J8-A3YH | Thyroid Carcinoma | Primary solid Tumor | C73 | stage i | 14470 | alive | NA | male | 1973 | white |
| TCGA-J8-A3YH-06A | TCGA-J8-A3YH | Thyroid Carcinoma | Metastatic | C73 | stage i | 14470 | alive | NA | male | 1973 | white |
| TCGA-J8-A4HW-01A | TCGA-J8-A4HW | Thyroid Carcinoma | Primary solid Tumor | C73 | stage iii | 21587 | alive | NA | female | 1953 | white |

| | | | | | | | | | | | |
|---|---|---|---|---|---|---|---|---|---|---|---|
| TCGA-J8-A4HW-06A | TCGA-J8-A4HW | Thyroid Carcinoma | Metastatic | C73 | stage iii | 21587 | alive | NA | female | 1953 | white |
| TCGA-KU-A6H7-01A | TCGA-KU-A6H7 | Head and Neck Squamous Cell Carcinoma | Primary solid Tumor | C09.9 | stage iva | 20314 | alive | NA | female | 1958 | white |
| TCGA-KU-A6H7-06A | TCGA-KU-A6H7 | Head and Neck Squamous Cell Carcinoma | Metastatic | C09.9 | stage iva | 20314 | alive | NA | female | 1958 | white |
| TCGA-NH-A8F7-01A | TCGA-NH-A8F7 | Colon Adenocarcinoma | Primary solid Tumor | C18.7 | stage iia | 19535 | alive | NA | female | 1960 | black or african american |
| TCGA-NH-A8F7-06A | TCGA-NH-A8F7 | Colon Adenocarcinoma | Metastatic | C18.7 | stage iia | 19535 | alive | NA | female | 1960 | black or african american |
| TCGA-RW-A686-01A | TCGA-RW-A686 | Pheochromocytoma and Paraganglioma | Primary solid Tumor | C74.9 | not reported | 22718 | alive | NA | male | 1950 | white |
| TCGA-RW-A686-06A | TCGA-RW-A686 | Pheochromocytoma and Paraganglioma | Metastatic | C74.9 | not reported | 22718 | alive | NA | male | 1950 | white |
| TCGA-SI-A71O-01A | TCGA-SI-A71O | Sarcoma | Primary solid Tumor | C49.2 | not reported | 10710 | dead | 694 | male | 1984 | white |
| TCGA-SI-A71O-06A | TCGA-SI-A71O | Sarcoma | Metastatic | C49.2 | not reported | 10710 | dead | 694 | male | 1984 | white |
| TCGA-SR-A6MX-01A | TCGA-SR-A6MX | Pheochromocytoma and Paraganglioma | Primary solid Tumor | C72.9 | not reported | 20186 | dead | 95 | male | 1958 | white |

| | | | | | | | | | | | |
|---|---|---|---|---|---|---|---|---|---|---|---|
| TCGA-SR-A6MX-06A | TCGA-SR-A6MX | Pheochromocytoma and Paraganglioma | Metastatic | C72.9 | not reported | 20186 | dead | 95 | male | 1958 | white |
| TCGA-UC-A7PG-01A | TCGA-UC-A7PG | Cervical Squamous Cell Carcinoma and Endocervical Adenocarcinoma | Primary solid Tumor | C53.9 | not reported | 16231 | dead | 370 | female | 1959 | white |
| TCGA-UC-A7PG-06A | TCGA-UC-A7PG | Cervical Squamous Cell Carcinoma and Endocervical Adenocarcinoma | Metastatic | C53.9 | not reported | 16231 | dead | 370 | female | 1959 | white |
| TCGA-UF-A71A-01A | TCGA-UF-A71A | Head and Neck Squamous Cell Carcinoma | Primary solid Tumor | C04.9 | stage iva | 24623 | dead | 86 | male | 1942 | white |
| TCGA-UF-A71A-06A | TCGA-UF-A71A | Head and Neck Squamous Cell Carcinoma | Metastatic | C04.9 | stage iva | 24623 | dead | 86 | male | 1942 | white |
| TCGA-V1-A9O5-01A | TCGA-V1-A9O5 | Prostate Adenocarcinoma | Primary solid Tumor | C61 | not reported | 23725 | alive | NA | male | NA | not reported |

| | | | | | | | | | | | |
|---|---|---|---|---|---|---|---|---|---|---|---|
| TCGA-V1-A9O5-06A | TCGA-V1-A9O5 | Prostate Adenocarcinoma | Metastatic | C61 | not reported | 23725 | alive | NA | male | NA | not reported |
| TCGA-V5-A7RC-01B | TCGA-V5-A7RC | Esophageal Carcinoma | Primary solid Tumor | C15.4 | not reported | 20439 | dead | 104 | male | NA | black or african american |
| TCGA-V5-A7RC-06A | TCGA-V5-A7RC | Esophageal Carcinoma | Metastatic | C15.4 | not reported | 20439 | dead | 104 | male | NA | black or african american |

| Edge importance | miRNA transcript | mRNA gene |
|---|---|---|
| 0.5327944 | hsa.mir.450a.1 | ENSG00000134363 |
| 0.5149385 | hsa.mir.452 | ENSG00000156103 |
| 0.4582074 | hsa.mir.1.2 | ENSG00000137809 |
| 0.5016520 | hsa.mir.27b | ENSG00000157551 |
| 0.4467838 | hsa.mir.374b | ENSG00000142661 |
| 0.4788752 | hsa.mir.374b | ENSG00000171812 |
| 0.5259402 | hsa.mir.3117 | ENSG00000164107 |
| 0.4910721 | hsa.mir.301a | ENSG00000174099 |
| 0.4247100 | hsa.mir.570 | ENSG00000204262 |
| 0.4383128 | hsa.mir.450a.2 | ENSG00000176971 |
| 0.4236204 | hsa.mir.3912 | ENSG00000166923 |
| 0.4766608 | hsa.mir.3912 | ENSG00000227039 |
| 0.4249912 | hsa.mir.374a | ENSG00000180044 |
| 0.5095255 | hsa.mir.338 | ENSG00000232679 |
| 0.4384886 | hsa.mir.22 | ENSG00000180448 |
| 0.4766257 | hsa.mir.1.1 | ENSG00000134363 |
| 0.5006678 | hsa.mir.450b | ENSG00000075035 |
| 0.4233392 | hsa.mir.450b | ENSG00000156103 |
| 0.5413005 | hsa.mir.424 | ENSG00000137809 |
| 0.4422847 | hsa.mir.542 | ENSG00000157551 |
| 0.4262917 | hsa.mir.542 | ENSG00000189320 |
| 0.5082953 | hsa.mir.143 | ENSG00000142661 |
| 0.4728295 | hsa.mir.548b | ENSG00000164107 |
| 0.4434446 | hsa.mir.2355 | ENSG00000174099 |
| 0.4763796 | hsa.mir.592 | ENSG00000165617 |
| 0.4213357 | hsa.mir.592 | ENSG00000204262 |
| 0.4594376 | hsa.mir.4677 | ENSG00000176971 |

| | | |
|---|---|---|
| 0.5599297 | hsa.mir.590 | ENSG00000166923 |
| 0.4647100 | hsa.mir.454 | ENSG00000145824 |
| 0.5232337 | hsa.mir.454 | ENSG00000180044 |
| 0.4897715 | hsa.mir.135a.1 | ENSG00000167483 |
| 0.4387698 | hsa.mir.660 | ENSG00000156103 |
| 0.4711775 | hsa.mir.660 | ENSG00000180448 |
| 0.4401757 | hsa.mir.1277 | ENSG00000075035 |
| 0.5049561 | hsa.mir.1277 | ENSG00000186081 |
| 0.4619684 | hsa.mir.190a | ENSG00000137809 |
| 0.4985589 | hsa.mir.190a | ENSG00000171812 |
| 0.4977856 | hsa.mir.4999 | ENSG00000105989 |
| 0.4515641 | hsa.mir.4999 | ENSG00000189320 |
| 0.4577153 | hsa.mir.32 | ENSG00000142661 |
| 0.4479789 | hsa.mir.185 | ENSG00000204262 |
| 0.5437610 | hsa.mir.1270 | ENSG00000145244 |
| 0.4754306 | hsa.let.7c | ENSG00000165617 |
| 0.4803866 | hsa.let.7c | ENSG00000227039 |
| 0.5892091 | hsa.mir.26a.1 | ENSG00000145423 |
| 0.4546924 | hsa.mir.653 | ENSG00000133110 |
| 0.5261863 | hsa.mir.653 | ENSG00000166923 |
| 0.4245343 | hsa.mir.580 | ENSG00000145824 |
| 0.4788049 | hsa.mir.580 | ENSG00000180044 |
| 0.5224253 | hsa.mir.3065 | ENSG00000167483 |
| 0.4653427 | hsa.let.7i | ENSG00000156103 |
| 0.5205624 | hsa.let.7i | ENSG00000180448 |
| 0.4229525 | hsa.mir.26a.2 | ENSG00000137809 |
| 0.4330756 | hsa.mir.26a.2 | ENSG00000168071 |
| 0.4721265 | hsa.mir.3926.2 | ENSG00000157551 |
| 0.4669244 | hsa.mir.3926.2 | ENSG00000186081 |

| | | |
|---|---|---|
| 0.4910721 | hsa.mir.126 | ENSG00000171812 |
| 0.4653076 | hsa.mir.301b | ENSG00000105989 |
| 0.4659051 | hsa.mir.301b | ENSG00000156103 |
| 0.4360633 | hsa.mir.301b | ENSG00000164107 |
| 0.4333568 | hsa.mir.301b | ENSG00000189320 |
| 0.4287170 | hsa.mir.95 | ENSG00000142661 |
| 0.5902988 | hsa.mir.101.2 | ENSG00000110427 |
| 0.4507909 | hsa.mir.101.2 | ENSG00000157551 |
| 0.4289279 | hsa.mir.101.2 | ENSG00000204262 |
| 0.4147979 | hsa.mir.204 | ENSG00000171812 |
| 0.4892091 | hsa.mir.101.1 | ENSG00000122986 |
| 0.4328647 | hsa.mir.130a | ENSG00000145423 |
| 0.4273462 | hsa.mir.130a | ENSG00000174099 |
| 0.4664323 | hsa.mir.891a | ENSG00000232679 |
| 0.5379613 | hsa.mir.19a | ENSG00000145824 |
| 0.4522671 | hsa.mir.98 | ENSG00000134363 |
| 0.4403866 | hsa.mir.98 | ENSG00000227039 |
| 0.4491740 | hsa.mir.4797 | ENSG00000156103 |
| 0.4208787 | hsa.mir.655 | ENSG00000137809 |
| 0.4293497 | hsa.mir.655 | ENSG00000168071 |
| 0.4404921 | hsa.mir.449a | ENSG00000157551 |
| 0.4447803 | hsa.mir.449a | ENSG00000186081 |
| 0.4382777 | hsa.mir.152 | ENSG00000142661 |
| 0.4309666 | hsa.mir.152 | ENSG00000171812 |
| 0.4604218 | hsa.mir.552 | ENSG00000164107 |
| 0.4811248 | hsa.mir.552 | ENSG00000189320 |
| 0.4312127 | hsa.mir.708 | ENSG00000171812 |
| 0.4730404 | hsa.mir.708 | ENSG00000174099 |
| 0.4478735 | hsa.mir.1228 | ENSG00000165617 |

| | | |
|---|---|---|
| 0.5303691 | hsa.mir.1228 | ENSG00000204262 |
| 0.4171178 | hsa.mir.9.3 | ENSG00000166923 |
| 0.5064323 | hsa.mir.9.3 | ENSG00000227039 |
| 0.4395782 | hsa.mir.9.2 | ENSG00000180044 |
| 0.4272056 | hsa.mir.29c | ENSG00000133110 |
| 0.4190861 | hsa.mir.421 | ENSG00000137809 |
| 0.4157469 | hsa.mir.1179 | ENSG00000134363 |
| 0.4285413 | hsa.mir.153.2 | ENSG00000165617 |
| 0.4410896 | hsa.mir.376c | ENSG00000176971 |
| 0.4365554 | hsa.mir.135a.2 | ENSG00000176971 |
| 0.4695255 | hsa.mir.217 | ENSG00000166923 |
| 0.4225308 | hsa.let.7g | ENSG00000180044 |
| 0.4511424 | hsa.mir.16.2 | ENSG00000133110 |
| 0.4201406 | hsa.mir.18a | ENSG00000105989 |
| 0.4388049 | hsa.mir.100 | ENSG00000189320 |
| 0.4444288 | hsa.mir.5690 | ENSG00000142661 |
| 0.4597540 | hsa.mir.5690 | ENSG00000165617 |
| 0.4325483 | hsa.mir.6868 | ENSG00000204262 |
| 0.4701230 | hsa.mir.6815 | ENSG00000176971 |
| 0.4265026 | hsa.mir.216a | ENSG00000171812 |
| 0.4175747 | hsa.mir.543 | ENSG00000174099 |
| 0.4356415 | hsa.mir.26b | ENSG00000204262 |
| 0.4857996 | hsa.mir.29a | ENSG00000142661 |
| 0.5115290 | hsa.mir.7.3 | ENSG00000110427 |
| 0.4462566 | hsa.mir.4444.1 | ENSG00000133110 |
| 0.4577153 | hsa.mir.20a | ENSG00000122986 |
| 0.5384886 | hsa.mir.212 | ENSG00000145423 |
| 0.4974692 | hsa.mir.451a | ENSG00000145824 |
| 0.4865729 | hsa.mir.451a | ENSG00000180044 |

| | | |
|---|---|---|
| 0.4837610 | hsa.mir.548o.2 | ENSG00000075035 |
| 0.5343409 | hsa.mir.548o.2 | ENSG00000167483 |
| 0.4286819 | hsa.mir.548o.2 | ENSG00000186081 |
| 0.4302636 | hsa.mir.3684 | ENSG00000133110 |
| 0.4581019 | hsa.mir.3684 | ENSG00000164107 |
| 0.4400351 | hsa.mir.3684 | ENSG00000168071 |
| 0.4337786 | hsa.mir.514a.3 | ENSG00000075035 |
| 0.4192970 | hsa.mir.514a.3 | ENSG00000157551 |
| 0.4743409 | hsa.mir.30e | ENSG00000105989 |
| 0.4733568 | hsa.mir.30e | ENSG00000189320 |
| 0.4332865 | hsa.mir.154 | ENSG00000227039 |
| 0.4768014 | hsa.mir.506 | ENSG00000122986 |
| 0.4686116 | hsa.mir.29b.2 | ENSG00000145824 |
| 0.4366608 | hsa.mir.299 | ENSG00000156103 |
| 0.4597540 | hsa.mir.17 | ENSG00000168071 |
| 0.4214060 | hsa.mir.4636 | ENSG00000157551 |
| 0.4461863 | hsa.mir.5696 | ENSG00000171812 |
| 0.4349385 | hsa.mir.19b.1 | ENSG00000137809 |
| 0.4394728 | hsa.mir.19b.1 | ENSG00000174099 |
| 0.5044991 | hsa.mir.598 | ENSG00000204262 |
| 0.4404569 | hsa.mir.137 | ENSG00000122986 |
| 0.4416169 | hsa.mir.137 | ENSG00000227039 |
| 0.4572232 | hsa.mir.509.1 | ENSG00000133110 |
| 0.4350439 | hsa.mir.887 | ENSG00000180448 |
| 0.4190158 | hsa.mir.643 | ENSG00000075035 |
| 0.4465378 | hsa.mir.15a | ENSG00000168071 |
| 0.4834095 | hsa.mir.320c.2 | ENSG00000157551 |
| 0.4295958 | hsa.mir.1468 | ENSG00000171812 |
| 0.5037258 | hsa.mir.335 | ENSG00000164107 |

| | | |
|---|---|---|
| 0.4778559 | hsa.mir.133a.2 | ENSG00000174099 |
| 0.4373286 | hsa.mir.618 | ENSG00000165617 |
| 0.4382425 | hsa.mir.33b | ENSG00000176971 |
| 0.4480141 | hsa.mir.138.2 | ENSG00000227039 |
| 0.4289279 | hsa.mir.509.2 | ENSG00000180044 |
| 0.4739895 | hsa.mir.1251 | ENSG00000134363 |
| 0.4661160 | hsa.mir.1245a | ENSG00000075035 |
| 0.4617575 | hsa.mir.4676 | ENSG00000137809 |
| 0.4248155 | hsa.mir.196b | ENSG00000157551 |
| 0.5078735 | hsa.mir.6516 | ENSG00000142661 |
| 0.4714587 | hsa.let.7f.1 | ENSG00000164107 |
| 0.4719508 | hsa.mir.4786 | ENSG00000166923 |
| 0.4153251 | hsa.mir.4786 | ENSG00000176971 |
| 0.4653779 | hsa.mir.6125 | ENSG00000167483 |
| 0.4618981 | hsa.mir.6125 | ENSG00000232679 |
| 0.4156415 | hsa.mir.1343 | ENSG00000180448 |
| 0.4184183 | hsa.mir.188 | ENSG00000168071 |
| 0.4898418 | hsa.mir.218.1 | ENSG00000075035 |
| 0.4691037 | hsa.mir.30c.1 | ENSG00000137809 |
| 0.4328998 | hsa.mir.24.1 | ENSG00000189320 |
| 0.5060457 | hsa.mir.146b | ENSG00000145244 |
| 0.4842179 | hsa.mir.3157 | ENSG00000145423 |
| 0.4207030 | hsa.mir.3157 | ENSG00000176971 |
| 0.4194728 | hsa.mir.153.1 | ENSG00000166923 |
| 0.4171529 | hsa.mir.376b | ENSG00000180044 |
| 0.4168717 | hsa.mir.136 | ENSG00000156103 |
| 0.6023902 | hsa.mir.136 | ENSG00000180448 |
| 0.4200000 | hsa.mir.3199.1 | ENSG00000133110 |
| 0.4300879 | hsa.mir.3199.1 | ENSG00000168071 |

| | | |
|---|---|---|
| 0.4213005 | hsa.mir.581 | ENSG00000075035 |
| 0.4390861 | hsa.mir.34c | ENSG00000176971 |
| 0.4171178 | hsa.mir.372 | ENSG00000142661 |
| 0.4650615 | hsa.mir.495 | ENSG00000145824 |
| 0.4423902 | hsa.mir.889 | ENSG00000134363 |
| 0.4893146 | hsa.mir.889 | ENSG00000166923 |
| 0.4200351 | hsa.mir.145 | ENSG00000105989 |
| 0.4652021 | hsa.mir.5706 | ENSG00000145824 |
| 0.4808436 | hsa.mir.378d.1 | ENSG00000134363 |
| 0.4229525 | hsa.mir.5001 | ENSG00000105989 |
| 0.5171529 | hsa.mir.500b | ENSG00000156103 |
| 0.4194025 | hsa.mir.551a | ENSG00000168071 |
| 0.4360984 | hsa.mir.6875 | ENSG00000174099 |
| 0.4448506 | hsa.mir.1296 | ENSG00000110427 |
| 0.4243234 | hsa.mir.1296 | ENSG00000189320 |
| 0.4245343 | hsa.mir.1296 | ENSG00000204262 |
| 0.4437961 | hsa.mir.504 | ENSG00000122986 |
| 0.4173638 | hsa.mir.486.1 | ENSG00000134363 |
| 0.4408787 | hsa.mir.1180 | ENSG00000232679 |
| 0.4438313 | hsa.mir.491 | ENSG00000145423 |
| 0.4756415 | hsa.mir.491 | ENSG00000168071 |
| 0.4381019 | hsa.mir.1248 | ENSG00000075035 |
| 0.4250967 | hsa.mir.556 | ENSG00000189320 |
| 0.4446397 | hsa.mir.576 | ENSG00000145244 |
| 0.4812302 | hsa.let.7a.3 | ENSG00000232679 |
| 0.4334271 | hsa.let.7a.1 | ENSG00000166923 |
| 0.4198946 | hsa.let.7a.1 | ENSG00000180448 |
| 0.5041828 | hsa.mir.181b.2 | ENSG00000157551 |
| 0.5036907 | hsa.mir.181a.2 | ENSG00000142661 |

| | | |
|---|---|---|
| 0.4394376 | hsa.mir.20b | ENSG00000110427 |
| 0.4928647 | hsa.mir.93 | ENSG00000227039 |
| 0.5111424 | hsa.mir.550a.1 | ENSG00000176971 |
| 0.4661511 | hsa.mir.6503 | ENSG00000133110 |
| 0.4744815 | hsa.mir.6503 | ENSG00000232679 |
| 0.4200351 | hsa.mir.1247 | ENSG00000137809 |
| 0.4481547 | hsa.mir.410 | ENSG00000137809 |
| 0.4277329 | hsa.mir.432 | ENSG00000189320 |
| 0.4667135 | hsa.mir.129.1 | ENSG00000174099 |
| 0.4373989 | hsa.mir.4521 | ENSG00000156103 |
| 0.4503691 | hsa.mir.3614 | ENSG00000227039 |
| 0.5087170 | hsa.mir.379 | ENSG00000145423 |
| 0.4746573 | hsa.mir.379 | ENSG00000145824 |
| 0.4691740 | hsa.mir.4473 | ENSG00000145824 |
| 0.4349033 | hsa.mir.4473 | ENSG00000156103 |
| 0.4234095 | hsa.mir.6874 | ENSG00000133110 |
| 0.4159930 | hsa.mir.1254.2 | ENSG00000204262 |
| 0.4228120 | hsa.mir.324 | ENSG00000232679 |
| 0.4586643 | hsa.mir.4707 | ENSG00000180044 |
| 0.4431283 | hsa.mir.433 | ENSG00000232679 |
| 0.4543761 | hsa.mir.6833 | ENSG00000137809 |
| 0.4238664 | hsa.mir.3199.2 | ENSG00000165617 |
| 0.4236555 | hsa.mir.6510 | ENSG00000180448 |
| 0.4474517 | hsa.mir.425 | ENSG00000075035 |
| 0.4292091 | hsa.mir.4683 | ENSG00000142661 |
| 0.4186995 | hsa.mir.193a | ENSG00000189320 |
| 0.4159930 | hsa.mir.30b | ENSG00000176971 |
| 0.4165554 | hsa.mir.1305 | ENSG00000180448 |
| 0.4461160 | hsa.mir.7854 | ENSG00000168071 |

| | | |
|---|---|---|
| 0.4298418 | hsa.mir.636 | ENSG00000145824 |
| 0.4288225 | hsa.mir.636 | ENSG00000174099 |
| 0.4447803 | hsa.mir.320b.2 | ENSG00000133110 |
| 0.4618278 | hsa.mir.320b.2 | ENSG00000204262 |
| 0.4655888 | hsa.mir.378c | ENSG00000227039 |
| 0.4469244 | hsa.mir.5586 | ENSG00000137809 |
| 0.5018278 | hsa.mir.934 | ENSG00000156103 |
| 0.4395782 | hsa.mir.1229 | ENSG00000227039 |
| 0.4342004 | hsa.mir.4687 | ENSG00000122986 |
| 0.4373286 | hsa.mir.4687 | ENSG00000166923 |
| 0.4172935 | hsa.mir.4687 | ENSG00000227039 |
| 0.4228822 | hsa.mir.496 | ENSG00000145423 |
| 0.4228822 | hsa.mir.31 | ENSG00000232679 |
| 0.4241476 | hsa.mir.3187 | ENSG00000171812 |
| 0.4653427 | hsa.mir.632 | ENSG00000164107 |
| 0.4313181 | hsa.mir.190b | ENSG00000145244 |
| 0.4183128 | hsa.mir.3934 | ENSG00000133110 |
| 0.4447100 | hsa.mir.3934 | ENSG00000167483 |
| 0.4160281 | hsa.mir.7.2 | ENSG00000105989 |
| 0.4671002 | hsa.mir.652 | ENSG00000145244 |
| 0.4959930 | hsa.mir.296 | ENSG00000165617 |
| 0.4199649 | hsa.mir.365a | ENSG00000204262 |
| 0.4338489 | hsa.mir.3940 | ENSG00000180044 |
| 0.5054130 | hsa.mir.181a.1 | ENSG00000167483 |
| 0.4218278 | hsa.mir.5698 | ENSG00000180448 |
| 0.4343058 | hsa.mir.6837 | ENSG00000157551 |
| 0.4253076 | hsa.mir.4461 | ENSG00000145244 |
| 0.4501933 | hsa.mir.1291 | ENSG00000122986 |
| 0.4219332 | hsa.mir.4449 | ENSG00000145423 |

| | | |
|---|---|---|
| 0.4431283 | hsa.mir.3651 | ENSG00000174099 |
| 0.5036555 | hsa.mir.4762 | ENSG00000168071 |
| 0.4267487 | hsa.mir.223 | ENSG00000157551 |
| 0.5311424 | hsa.mir.223 | ENSG00000186081 |
| 0.5043585 | hsa.mir.15b | ENSG00000171812 |
| 0.5707909 | hsa.mir.874 | ENSG00000189320 |
| 0.4198946 | hsa.mir.1292 | ENSG00000137809 |
| 0.4682953 | hsa.mir.1292 | ENSG00000174099 |
| 0.4409139 | hsa.mir.30d | ENSG00000110427 |
| 0.5165905 | hsa.mir.30d | ENSG00000204262 |
| 0.4500879 | hsa.mir.579 | ENSG00000122986 |
| 0.4297012 | hsa.mir.579 | ENSG00000164107 |
| 0.4709666 | hsa.mir.3130.1 | ENSG00000133110 |
| 0.4754657 | hsa.mir.3130.1 | ENSG00000232679 |
| 0.4666432 | hsa.mir.769 | ENSG00000145824 |
| 0.4585940 | hsa.mir.744 | ENSG00000075035 |
| 0.4918805 | hsa.mir.744 | ENSG00000156103 |
| 0.4739192 | hsa.mir.5090 | ENSG00000137809 |
| 0.4928998 | hsa.mir.5090 | ENSG00000168071 |
| 0.4791916 | hsa.mir.939 | ENSG00000157551 |
| 0.4742355 | hsa.mir.4668 | ENSG00000171812 |
| 0.4676977 | hsa.mir.345 | ENSG00000164107 |
| 0.4170475 | hsa.mir.345 | ENSG00000189320 |
| 0.4434798 | hsa.mir.3170 | ENSG00000174099 |
| 0.4444288 | hsa.mir.1306 | ENSG00000204262 |
| 0.4619332 | hsa.mir.128.1 | ENSG00000176971 |
| 0.4335677 | hsa.mir.4444.2 | ENSG00000122986 |
| 0.4203866 | hsa.mir.4444.2 | ENSG00000166923 |
| 0.5365202 | hsa.mir.4444.2 | ENSG00000227039 |

| | | |
|---|---|---|
| 0.4337786 | hsa.mir.128.2 | ENSG00000145824 |
| 0.4727944 | hsa.mir.128.2 | ENSG00000180044 |
| 0.4559578 | hsa.mir.4484 | ENSG00000133110 |
| 0.4315290 | hsa.mir.4484 | ENSG00000167483 |
| 0.5444640 | hsa.mir.4484 | ENSG00000232679 |
| 0.4843585 | hsa.mir.4781 | ENSG00000180448 |
| 0.5469244 | hsa.mir.3150b | ENSG00000134363 |
| 0.5003866 | hsa.mir.664a | ENSG00000075035 |
| 0.4645343 | hsa.mir.6820 | ENSG00000137809 |
| 0.4315993 | hsa.mir.501 | ENSG00000105989 |
| 0.4497364 | hsa.mir.1269a | ENSG00000142661 |
| 0.4643234 | hsa.mir.3178 | ENSG00000110427 |
| 0.4589104 | hsa.mir.3178 | ENSG00000164107 |
| 0.4530404 | hsa.mir.192 | ENSG00000145244 |
| 0.4953603 | hsa.mir.885 | ENSG00000165617 |
| 0.4464323 | hsa.mir.624 | ENSG00000145423 |
| 0.5504745 | hsa.mir.624 | ENSG00000176971 |
| 0.4525835 | hsa.mir.3607 | ENSG00000166923 |
| 0.4208084 | hsa.mir.4662a | ENSG00000145824 |
| 0.4851318 | hsa.mir.4662a | ENSG00000180044 |
| 0.4231986 | hsa.mir.138.1 | ENSG00000133110 |
| 0.4457996 | hsa.mir.3652 | ENSG00000180448 |
| 0.4477680 | hsa.mir.342 | ENSG00000168071 |
| 0.4999649 | hsa.mir.146a | ENSG00000075035 |
| 0.4469947 | hsa.mir.146a | ENSG00000186081 |
| 0.4949736 | hsa.mir.4661 | ENSG00000137809 |
| 0.4539895 | hsa.mir.4423 | ENSG00000189320 |
| 0.5226714 | hsa.mir.196a.2 | ENSG00000145244 |
| 0.5256591 | hsa.mir.5684 | ENSG00000145423 |

| | | |
|---|---|---|
| 0.4620387 | hsa.mir.532 | ENSG00000167483 |
| 0.4429525 | hsa.mir.139 | ENSG00000180448 |
| 0.4499824 | hsa.mir.6514 | ENSG00000157551 |
| 0.5482953 | hsa.mir.378d.2 | ENSG00000189320 |
| 0.4434446 | hsa.mir.584 | ENSG00000110427 |
| 0.4451318 | hsa.mir.584 | ENSG00000145423 |
| 0.4473111 | hsa.mir.4791 | ENSG00000168071 |
| 0.5277680 | hsa.mir.6842 | ENSG00000145824 |
| 0.4484007 | hsa.mir.6842 | ENSG00000168071 |
| 0.4416520 | hsa.mir.3928 | ENSG00000156103 |
| 0.4572232 | hsa.mir.1226 | ENSG00000167483 |
| 0.4161336 | hsa.mir.5091 | ENSG00000156103 |
| 0.4416169 | hsa.mir.92a.2 | ENSG00000186081 |
| 0.4906151 | hsa.mir.92a.1 | ENSG00000189320 |
| 0.5149385 | hsa.mir.194.1 | ENSG00000110427 |
| 0.4170475 | hsa.mir.4664 | ENSG00000145244 |
| 0.4449209 | hsa.mir.4724 | ENSG00000122986 |
| 0.4512127 | hsa.mir.3193 | ENSG00000133110 |
| 0.5057645 | hsa.mir.1976 | ENSG00000145824 |
| 0.4297012 | hsa.mir.151b | ENSG00000180448 |
| 0.4533919 | hsa.mir.423 | ENSG00000137809 |
| 0.4181371 | hsa.mir.423 | ENSG00000156103 |
| 0.4376801 | hsa.mir.423 | ENSG00000168071 |
| 0.4278383 | hsa.mir.550a.2 | ENSG00000157551 |
| 0.4973638 | hsa.mir.3922 | ENSG00000171812 |
| 0.5113884 | hsa.mir.92b | ENSG00000204262 |
| 0.4802460 | hsa.mir.191 | ENSG00000227039 |
| 0.5135325 | hsa.mir.4491 | ENSG00000133110 |
| 0.4250264 | hsa.mir.4491 | ENSG00000232679 |

| | | |
|---|---|---|
| 0.4678735 | hsa.mir.1275 | ENSG00000145824 |
| 0.4211951 | hsa.mir.1275 | ENSG00000180448 |
| 0.5630580 | hsa.mir.135b | ENSG00000134363 |
| 0.4441828 | hsa.mir.3074 | ENSG00000075035 |
| 0.5090685 | hsa.mir.3074 | ENSG00000156103 |
| 0.4822847 | hsa.mir.194.2 | ENSG00000137809 |
| 0.5103691 | hsa.mir.6806 | ENSG00000157551 |
| 0.4746221 | hsa.mir.3127 | ENSG00000171812 |
| 0.4966960 | hsa.mir.203b | ENSG00000164107 |
| 0.4541301 | hsa.mir.659 | ENSG00000145244 |
| 0.4953954 | hsa.mir.659 | ENSG00000174099 |
| 0.4514587 | hsa.mir.6843 | ENSG00000165617 |
| 0.4954306 | hsa.mir.942 | ENSG00000176971 |
| 0.4283304 | hsa.mir.196a.1 | ENSG00000232679 |
| 0.4230580 | hsa.mir.3690.1 | ENSG00000145824 |
| 0.4164851 | hsa.mir.6716 | ENSG00000134363 |
| 0.4996134 | hsa.mir.10b | ENSG00000075035 |
| 0.4640070 | hsa.mir.1181 | ENSG00000137809 |
| 0.4568366 | hsa.mir.330 | ENSG00000157551 |
| 0.4343761 | hsa.mir.346 | ENSG00000142661 |
| 0.4861863 | hsa.mir.6511b.1 | ENSG00000164107 |
| 0.4213357 | hsa.mir.6511b.1 | ENSG00000186081 |
| 0.4294552 | hsa.mir.675 | ENSG00000145244 |
| 0.4490685 | hsa.mir.28 | ENSG00000176971 |
| 0.4810545 | hsa.mir.877 | ENSG00000180044 |
| 0.4224956 | hsa.mir.6852 | ENSG00000167483 |
| 0.4597540 | hsa.mir.6852 | ENSG00000232679 |
| 0.4551845 | hsa.mir.487a | ENSG00000180448 |
| 0.4681547 | hsa.mir.1284 | ENSG00000075035 |

| | | |
|---|---|---|
| 0.4411248 | hsa.mir.1284 | ENSG00000186081 |
| 0.4641828 | hsa.mir.767 | ENSG00000156103 |
| 0.4695255 | hsa.let.7b | ENSG00000105989 |
| 0.4208787 | hsa.mir.760 | ENSG00000204262 |
| 0.4951494 | hsa.mir.5703 | ENSG00000145244 |
| 0.4376098 | hsa.mir.1224 | ENSG00000165617 |
| 0.4207733 | hsa.mir.99b | ENSG00000133110 |
| 0.4292794 | hsa.mir.3677 | ENSG00000168071 |
| 0.4223902 | hsa.mir.10a | ENSG00000134363 |
| 0.4559578 | hsa.mir.625 | ENSG00000105989 |
| 0.5091037 | hsa.mir.625 | ENSG00000189320 |
| 0.4426362 | hsa.mir.1249 | ENSG00000110427 |
| 0.4282601 | hsa.mir.1249 | ENSG00000164107 |
| 0.4607733 | hsa.mir.1266 | ENSG00000133110 |
| 0.4559930 | hsa.mir.4638 | ENSG00000180044 |
| 0.4958875 | hsa.mir.548o | ENSG00000145824 |
| 0.4159227 | hsa.mir.6854 | ENSG00000167483 |
| 0.4381019 | hsa.mir.6511b.2 | ENSG00000157551 |
| 0.4157469 | hsa.mir.375 | ENSG00000186081 |
| 0.4300176 | hsa.mir.6877 | ENSG00000171812 |
| 0.4654833 | hsa.mir.326 | ENSG00000164107 |
| 0.4478383 | hsa.mir.3136 | ENSG00000227039 |
| 0.4640422 | hsa.mir.937 | ENSG00000122986 |
| 0.4534974 | hsa.mir.6720 | ENSG00000180044 |

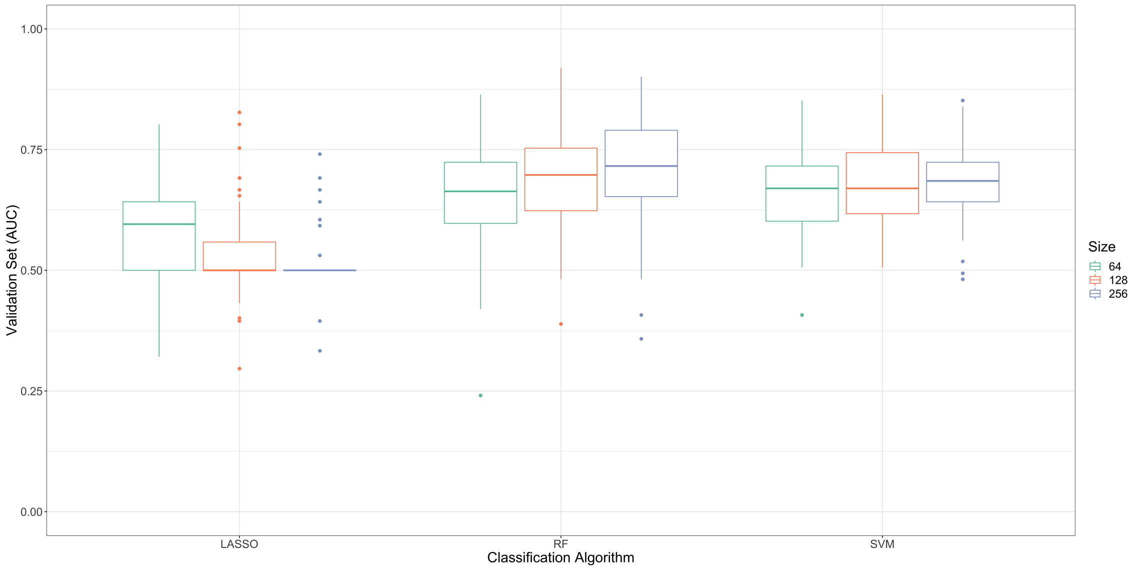


Figure 1: This figure shows the classification accuracy of metastatic status for paired tumour samples across 100 Monte-Carlo re-samplings (y-axis) for each classification model (x-axis) using the combined mRNA and miRNA expression. Boxplot colour refers to the top $N = \{64, 128, 256\}$ features as selected by Student's t-test. Although the classifier size somewhat impacts performance, LASSO is always outperformed by random forest (RF) and support vector machine (SVM) regardless of classifier size. Combining mRNA and miRNA biomarkers does not markedly improve performance over using only mRNA biomarkers.
