## Supplementary material for "A cross-cancer metastasis signature in the microRNA-mRNA axis of paired tissue samples"

```
setwd("/Volumes/all-members/genetics/TCGA")
library(magrittr)
library(stringr)

# Using tcga annotation files find patients with records
# for both primary and met tumours
annots <- list.files("annot", full.names = TRUE)
all <- lapply(annots, read.csv, stringsAsFactors = F)
master <- do.call(plyr::rbind.fill, all)
ms <- master[,!grepl("subtype", colnames(master))]
dupes <- names(table(ms$patient)[table(ms$patient) >= 2])
ms2 <- ms[ms$patient %in% dupes,]
ms2
table(ms2$definition)
pat.with.met <- ms2[ms2$definition == "Metastatic","patient"]
table(ms2[ms2$patient %in% pat.with.met, "definition"]) # 28 = metastatic +
primary solid
pat.with.pst <- ms2[ms2$definition == "Primary solid Tumor","patient"]
golden28 <- intersect(pat.with.met, pat.with.pst)
ms28 <- ms2[ms2$patient %in% golden28,]

sort(table(ms28[ms28$definition == "Primary solid Tumor", "name"]))
head(ms28)
write.csv(ms28, file = "golden28.csv")

####

# make golden 28 mRNA and miRNA expression matrices

g28 <- ms28[ms28$shortLetterCode %in% c("TP", "TM"),]
projects <- unique(g28$project_id)
for(type in c("tcga-miRNA", "tcga-ensg")){

  files <- lapply(
    projects,
    function(x){
      list.files(path = type, pattern = x, recursive = F, full.names = T)
    }
  )
  files <- unlist(files)


  raws <- lapply(
    files,
    read.csv,
    row.names = 1
  )

  ## check if all colnames are the same
  all(reduce(lapply(raws, rownames), intersect) == rownames(raws[[1]]))

  ## combine all df's
  allDF <- do.call("cbind", raws)

  ## fix col names to match sample name style
  colnames(allDF) <- colnames(allDF) %>%
    str_sub(end = 16) %>%
    str_replace_all("\\.", "-")

  # filter for golden28
  g28matrix <- allDF[, sort(intersect(golden28$sample, colnames(allDF)))]

  write.csv(
    g28matrix,
```

```
    file = file.path(outDir, paste(type, "golden28.csv", sep = '-'))
  )

}
```

```
# Import Golden 28 data
genes <- read.csv("Dropbox/Partners/Thin_Thom/wPaper/golden28/golden28-
genes.csv",
                  row.names = 1)
miRNA <- read.csv("Dropbox/Partners/Thin_Thom/wPaper/golden28/golden28-
mirna.csv",
                  row.names = 1)
annot <- read.csv("Dropbox/Partners/Thin_Thom/wPaper/golden28/golden28-
annot.csv")

# Missing data so Golden 28 becomes Golden 27
a55 <- colnames(miRNA)
a56 <- gsub("-", ".", annot[,2])
missingData <- annot[!a56 %in% a55 & annot$definition == "Metastatic",
"patient"]
removeSamples <- as.character(annot[annot$patient == missingData, "sample"])
removeSamples <- gsub("-", ".", removeSamples)
all(colnames(miRNA) %in% colnames(genes))
genes.27 <- genes[, !colnames(genes) %in% removeSamples]
miRNA.27 <- miRNA[, !colnames(miRNA) %in% removeSamples]
identical(colnames(genes.27), colnames(miRNA.27))
rownames(annot) <- a56
annot.27 <- annot[colnames(genes.27),]

# Save Golden 27 data
write.csv(genes.27, "Dropbox/Partners/Thin_Thom/wPaper/golden27/golden27-
genes.csv")
write.csv(miRNA.27, "Dropbox/Partners/Thin_Thom/wPaper/golden27/golden27-
mirna.csv")
write.csv(annot.27, "Dropbox/Partners/Thin_Thom/wPaper/golden27/golden27-
annot.csv")
```

```
# Import Golden 27 data
genes <- read.csv("golden27/golden27-genes.csv",
                  row.names = 1)
miRNA <- read.csv("golden27/golden27-mirna.csv",
                  row.names = 1)
annot <- read.csv("golden27/golden27-annot.csv")

# Count zeros
genes.0s <- apply(genes, 1, function(x) sum(x == 0) / ncol(genes))
hist(genes.0s)
miRNA.0s <- apply(miRNA, 1, function(x) sum(x == 0) / ncol(miRNA))
hist(miRNA.0s)
round(apply(apply(genes, 2, summary), 1, mean), 2)
round(apply(apply(miRNA, 2, summary), 1, mean), 2)

# Filter data
genes.half0s <- apply(genes, 1, function(x) sum(x == 0) > 27)
sum(genes.half0s)
miRNA.half0s <- apply(miRNA, 1, function(x) sum(x == 0) > 27)
sum(miRNA.half0s)
genes.sub <- genes[!genes.half0s,]
miRNA.sub <- miRNA[!miRNA.half0s,]

# Run DESeq2 on mRNA
library(DESeq2)
annot$definition <- factor(gsub(" ", "", annot$definition))
dds.g <- DESeqDataSetFromMatrix(genes.sub, annot, design = ~ patient +
definition)# + primary_site)
dds.g <- DESeq(dds.g)
res.g <- results(dds.g, contrast = c("definition", "PrimarysolidTumor",
"Metastatic"))

# Visualize results
plotMA(res.g)
vsd.g <- vst(dds.g, blind = FALSE)
plotPCA(vsd.g, intgroup = c("primary_site"))
plotPCA(vsd.g, intgroup = c("definition"))

# Run DEseq2 on miRNA
library(DESeq2)
#annot$definition <- factor(gsub(" ", "", annot$definition))
dds.m <- DESeqDataSetFromMatrix(miRNA.sub, annot, design = ~ patient +
definition)# + primary_site)
dds.m <- DESeq(dds.m)
res.m <- results(dds.m, contrast = c("definition", "PrimarysolidTumor",
"Metastatic"))

# Visualize results
plotMA(res.m)
vsd.m <- vst(dds.m, blind = FALSE, nsub = 200)
plotPCA(vsd.m, intgroup = c("primary_site"))
plotPCA(vsd.m, intgroup = c("definition"))

# Save normalized counts
write.csv(counts(dds.g, normalized = TRUE), file = "golden27/golden27-genes-
normal.csv")
write.csv(counts(dds.m, normalized = TRUE), file = "golden27/golden27-mirna-
normal.csv")
write.csv(annot, file = "golden27/golden27-annot-normal.csv")
write.csv(res.g, file = "2-DESeq2-genes.csv")
write.csv(res.m, file = "2-DESeq2-mirna.csv")
```

```
setwd("/home/thom/Dropbox/Partners/Thin_Thom/wPaper")

# Import Golden 27 data (normalized)
genes <- read.csv("golden27/golden27-genes-normal.csv",
                  row.names = 1)
genes <- t(genes)
miRNA <- read.csv("golden27/golden27-mirna-normal.csv",
                  row.names = 1)
miRNA <- t(miRNA)
annot <- read.csv("golden27/golden27-annot-normal.csv",
                  stringsAsFactors = FALSE)
identical(annot$X.1, rownames(miRNA))
identical(annot$X.1, rownames(genes))

de.genes <- read.csv("2-DESeq2-genes.csv", row.names = 1)
de.miRNA <- read.csv("2-DESeq2-mirna.csv", row.names = 1)

de.genes[is.na(de.genes$padj),] <- 1
de.genes <- de.genes[de.genes$padj < .05,]
de.miRNA[is.na(de.miRNA$padj),] <- 1
de.miRNA <- de.miRNA[de.miRNA$padj < .05,]

plot(de.genes$log2FoldChange)

# Recode gene expression as difference between primary and met
identical(
  annot[annot$definition == "PrimarysolidTumor", "patient"],
  annot[annot$definition == "Metastatic", "patient"]
)

genes <- genes + .5
genes.delta <- log2(genes[annot$definition == "Metastatic",] /
                      genes[annot$definition == "PrimarysolidTumor",])
delta.de <- genes.delta[,rownames(de.genes)]
min(delta.de)

dim(delta.de)
df <- propr::wide2long(delta.de)
head(df)
colnames(df) <- c("Delta", "Gene", "Patient")
cancers <- unique(annot[,c("sample", "disease_type")])
# cancers$disease_type <-
#   ifelse(!cancers$disease_type %in% c("Breast Invasive Carcinoma", "Thyroid
Carcinoma"),
#          "Other",
#          cancers$disease_type)
cancers$sample <- gsub("-", ".", cancers$sample)
df <- merge(df, cancers, by.x = "Patient", by.y = "sample")

library(ggplot2)
df[df$Gene %in% unique(df$Gene)[1:5],]
df$Gene <- factor(df$Gene, levels =
rownames(de.genes[order(de.genes$log2FoldChange),]))
df$disease_type <- factor(df$disease_type, levels =
names(sort(table(cancers$disease_type))))

jpeg("7-Fig1-viewDE.jpg",
     width = 20, height = 10, units = "in", res = 600)
ggplot(df,
       aes(x = Gene, y = Delta, group = disease_type, col = disease_type)) +
  geom_point(show.legend = FALSE, alpha = .4) + geom_smooth(method = "lm", se =
FALSE) +
  scale_color_brewer(palette = "Set3") +
  ylab("Log-fold change (Metastasis vs. Primary)") +
```

```
  xlab("Gene ID (Significant DE only)") +
  #ggtitle("Differential Expression (DE) of Pan-Cancer Metastasis Signals") +
  labs(col = "") +
  theme_bw() +
  scale_x_discrete(breaks = NULL) +
  theme(axis.text.x = element_text(angle = 90, hjust = 1, size = 0),
        text = element_text(size=18),
        plot.title = element_text(size=24),
        legend.position = "bottom")
dev.off()
```

```
splitPaired <- function(object, percent.include = 67, colBy, ...){

  # classCheck(object, "ExprsArray",
  #             "This function is applied to the results of ?exprso.")

  if(percent.include < 1 | percent.include > 100){
    stop("Uh oh! Use an inclusion percentage between 1-100!")
  }

  ids <- sample(unique(object@annot[,colBy]))

  # Return NULL validation set if percent.include = 100
  size <- round((length(ids) * percent.include)/100, digits = 0)
  if(size == ncol(object@exprs)){

    warning("splitSample built an empty validation set...\n\n")
    return(list(
      "array.train" = object,
      "array.valid" = NULL)
    )
  }

  random.train <- which(object@annot[,colBy] %in% ids[1:size])
  random.valid <- setdiff(1:nrow(object@annot), random.train)
  return(list(
    "array.train" = object[random.train, , drop = FALSE],
    "array.valid" = object[random.valid, , drop = FALSE])
  )
}

library(exprso)
setwd("/home/thom/Dropbox/Partners/Thin_Thom/wPaper/")

# Import Golden 27 data (normalized)
mRNA <- read.csv("golden27/golden27-genes-normal.csv",
                 row.names = 1)
mRNA <- t(mRNA)
annot <- read.csv("golden27/golden27-annot-normal.csv",
                  stringsAsFactors = FALSE)
identical(annot$X.1, rownames(mRNA))

# get the response variable
y <- annot[,c("definition", "disease_type", "patient")]

array <- exprso(x = mRNA, y = y)

ss <- splitPaired(array, colBy = "patient")
table(ss[[1]]@annot$patient)
table(ss[[2]]@annot$patient)

res <- lapply(c("buildLASSO", "buildRF", "buildSVM"), function(build){

  ss <- ctrlSplitSet(func = "splitPaired", colBy = "patient", percent.include =
67)
  fs <- ctrlFeatureSelect(func = "fsStats", top = 0, how = "t.test")
  gs <- ctrlGridSearch(func = "plGrid",
                       build,
                       top = c(64, 128, 256),
                       fold = NULL)

  plMonteCarlo(array, B = 100, ctrlSS = ss, ctrlFS = fs, ctrlGS = gs)
})

df <- do.call("conjoin", res)@summary
```

```
write.csv(df, file = "9a-mRNA-classifier.csv")
library(ggplot2)

df$build <- gsub("build", "", df$build)

jpeg("9a-Fig2-mRNA-classifier.jpg",
     width = 20, height = 10, units = "in", res = 600)
ggplot(df, aes(x = build, y = valid.auc, col = factor(top))) +
  geom_boxplot() + ylim(0, 1) + ylab("Validation Set (AUC)") +
  scale_color_brewer(palette = "Set2") +
  theme_bw() + xlab("Classification Algorithm") +
  labs(col = "Size") +
  theme(text = element_text(size=18))
dev.off()
```

```r
splitPaired <- function(object, percent.include = 67, colBy, ...){

  # classCheck(object, "ExprsArray",
  #            "This function is applied to the results of ?exprso.")

  if(percent.include < 1 | percent.include > 100){
    stop("Uh oh! Use an inclusion percentage between 1-100!")
  }

  ids <- sample(unique(object@annot[,colBy]))

  # Return NULL validation set if percent.include = 100
  size <- round((length(ids) * percent.include)/100, digits = 0)
  if(size == ncol(object@exprs)){

    warning("splitSample built an empty validation set...\n\n")
    return(list(
      "array.train" = object,
      "array.valid" = NULL)
    )
  }

  random.train <- which(object@annot[,colBy] %in% ids[1:size])
  random.valid <- setdiff(1:nrow(object@annot), random.train)
  return(list(
    "array.train" = object[random.train, , drop = FALSE],
    "array.valid" = object[random.valid, , drop = FALSE])
  )
}

library(exprso)
setwd("~/Dropbox/Thin_Thom/wPaper/")

# Import Golden 27 data (normalized)
miRNA <- read.csv("golden27/golden27-mirna-normal.csv",
                  row.names = 1)
miRNA <- t(miRNA)
annot <- read.csv("golden27/golden27-annot-normal.csv",
                  stringsAsFactors = FALSE)
identical(annot$X.1, rownames(miRNA))

# get the response variable
y <- annot[,c("definition", "disease_type", "patient")]

array <- exprso(x = miRNA, y = y)

ss <- splitPaired(array, colBy = "patient")
table(ss[[1]]@annot$patient)
table(ss[[2]]@annot$patient)

res <- lapply(c("buildLASSO", "buildRF", "buildSVM"), function(build){

  ss <- ctrlSplitSet(func = "splitPaired", colBy = "patient", percent.include =
67)
  fs <- ctrlFeatureSelect(func = "fsStats", top = 0, how = "t.test")
  gs <- ctrlGridSearch(func = "plGrid",
                       build,
                       top = c(64, 128, 256),
                       fold = NULL)

  plMonteCarlo(array, B = 100, ctrlSS = ss, ctrlFS = fs, ctrlGS = gs)
})

df <- do.call("conjoin", res)@summary
```

```
write.csv(df, file = "9b-miRNA-classifier.csv")
library(ggplot2)

df$build <- gsub("build", "", df$build)

jpeg("9b-Fig2-miRNA-classifier.jpg",
     width = 20, height = 10, units = "in", res = 600)
ggplot(df, aes(x = build, y = valid.auc, col = factor(top))) +
  geom_boxplot() + ylim(0, 1) + ylab("Validation Set (AUC)") +
  scale_color_brewer(palette = "Set2") +
  theme_bw() + xlab("Classification Algorithm") +
  labs(col = "Size") +
  theme(text = element_text(size=18))
dev.off()
```

```
splitPaired <- function(object, percent.include = 67, colBy, ...){

  # classCheck(object, "ExprsArray",
  #            "This function is applied to the results of ?exprso.")

  if(percent.include < 1 | percent.include > 100){
    stop("Uh oh! Use an inclusion percentage between 1-100!")
  }

  ids <- sample(unique(object@annot[,colBy]))

  # Return NULL validation set if percent.include = 100
  size <- round((length(ids) * percent.include)/100, digits = 0)
  if(size == ncol(object@exprs)){

    warning("splitSample built an empty validation set...\n\n")
    return(list(
      "array.train" = object,
      "array.valid" = NULL)
    )
  }

  random.train <- which(object@annot[,colBy] %in% ids[1:size])
  random.valid <- setdiff(1:nrow(object@annot), random.train)
  return(list(
    "array.train" = object[random.train, , drop = FALSE],
    "array.valid" = object[random.valid, , drop = FALSE])
  )
}

library(exprso)
# setwd("/home/thom/Dropbox/Partners/Thin_Thom/wPaper/")
setwd("~/Dropbox/Thin_Thom/wPaper")

# Import Golden 27 data (normalized)
mRNA <- read.csv("golden27/golden27-genes-normal.csv",
                 row.names = 1)
mRNA <- t(mRNA)

miRNA <- read.csv("golden27/golden27-mirna-normal.csv",
                  row.names = 1)
miRNA <- t(miRNA)

annot <- read.csv("golden27/golden27-annot-normal.csv",
                  stringsAsFactors = FALSE)
identical(annot$X.1, rownames(mRNA))
identical(annot$X.1, rownames(miRNA))
identical(rownames(mRNA), rownames(miRNA))

# get the response variable
y <- annot[,c("definition", "disease_type", "patient")]

array <- exprso(x = cbind(mRNA, miRNA), y = y)

ss <- splitPaired(array, colBy = "patient")
table(ss[[1]]@annot$patient)
table(ss[[2]]@annot$patient)

res <- lapply(c("buildLASSO", "buildRF", "buildSVM"), function(build){

  ss <- ctrlSplitSet(func = "splitPaired", colBy = "patient", percent.include =
67)
  fs <- ctrlFeatureSelect(func = "fsStats", top = 0, how = "t.test")
  gs <- ctrlGridSearch(func = "plGrid",
```

```
                       build,
                       top = c(64, 128, 256),
                       fold = NULL)

  plMonteCarlo(array, B = 100, ctrlSS = ss, ctrlFS = fs, ctrlGS = gs)
})

df <- do.call("conjoin", res)@summary
write.csv(df, file = "4c-mRNA-miRNA-classifier.csv")
library(ggplot2)

df$build <- gsub("build", "", df$build)

jpeg("4c-Fig2-mRNA-miRNA-classifier.jpg",
     width = 20, height = 10, units = "in", res = 600)
ggplot(df, aes(x = build, y = valid.auc, col = factor(top))) +
  geom_boxplot() + ylim(0, 1) + ylab("Validation Set (AUC)") +
  scale_color_brewer(palette = "Set2") +
  theme_bw() + xlab("Classification Algorithm") +
  labs(col = "Size") +
  theme(text = element_text(size=18))
dev.off()
```

```
setwd("/home/thom/Dropbox/Partners/Thin_Thom/wPaper")

# Import Golden 27 data (normalized)
genes <- read.csv("golden27/golden27-genes-normal.csv",
                  row.names = 1)
genes <- t(genes)
miRNA <- read.csv("golden27/golden27-mirna-normal.csv",
                  row.names = 1)
miRNA <- t(miRNA)
annot <- read.csv("golden27/golden27-annot-normal.csv",
                  stringsAsFactors = FALSE)
identical(annot$X.1, rownames(miRNA))
identical(annot$X.1, rownames(genes))

# Log transform
genes <- log(genes + 1)
miRNA <- log(miRNA + 1)

# Get DE genes only
de.genes <- read.csv("2-DESeq2-genes.csv", row.names = 1)
de.genes[is.na(de.genes$padj),] <- 1
de.genes <- de.genes[de.genes$padj < .05,]
genes <- genes[,rownames(de.genes)]

# Recode gene expression as difference between primary and met
identical(
  annot[annot$definition == "PrimarysolidTumor", "patient"],
  annot[annot$definition == "Metastatic", "patient"]
)
miRNA.delta <- miRNA[annot$definition == "PrimarysolidTumor",] -
  miRNA[annot$definition == "Metastatic",]

# Recode gene expression as difference between primary and met
identical(
  annot[annot$definition == "PrimarysolidTumor", "patient"],
  annot[annot$definition == "Metastatic", "patient"]
)
genes.delta <- genes[annot$definition == "PrimarysolidTumor",] -
  genes[annot$definition == "Metastatic",]

dim(genes.delta)
dim(miRNA.delta)

library(exprso)
set.seed(1)
how <- "buildRF"

res.null <- lapply(1:ncol(genes.delta), function(g){

  print(g)

  y.i <- genes.delta[,g]

  library(exprso)
  e.i <- exprso(miRNA.delta, y.i)
  ss <- ctrlSplitSet(func = "splitSample", percent.include = 67)
  fs <- ctrlFeatureSelect("fsCor")
  gs <- ctrlGridSearch(func = "plGrid", top = 128, how = how, fold = NULL)
  boot.i <- plMonteCarlo(e.i, B = 50, ctrlSS = ss, ctrlFS = fs, ctrlGS = gs)
  getWeights(boot.i)
})

# Name and save results
res.df <- lapply(1:length(res.null), function(r){
```

```
  data.frame("r" = colnames(genes.delta)[r], res.null[[r]])
})
res.df <- do.call(plyr::rbind.fill, res.df)
write.csv(res.df, file = paste0("5a-DeltaDelta-", how, "-LOG.csv"))
```

```
setwd("/home/thom/Dropbox/Partners/Thin_Thom/wPaper")

# Import Golden 27 data (normalized)
genes <- read.csv("golden27/golden27-genes-normal.csv",
                  row.names = 1)
genes <- t(genes)
miRNA <- read.csv("golden27/golden27-mirna-normal.csv",
                  row.names = 1)
miRNA <- t(miRNA)
annot <- read.csv("golden27/golden27-annot-normal.csv",
                  stringsAsFactors = FALSE)
identical(annot$X.1, rownames(miRNA))
identical(annot$X.1, rownames(genes))

# Log transform
genes <- log(genes + 1)
miRNA <- log(miRNA + 1)

# Get DE genes only
de.genes <- read.csv("2-DESeq2-genes.csv", row.names = 1)
de.genes[is.na(de.genes$padj),] <- 1
de.genes <- de.genes[de.genes$padj < .05,]
genes <- genes[,rownames(de.genes)]

# Recode gene expression as difference between primary and met
identical(
  annot[annot$definition == "PrimarysolidTumor", "patient"],
  annot[annot$definition == "Metastatic", "patient"]
)
miRNA.delta <- miRNA[annot$definition == "PrimarysolidTumor",] -
  miRNA[annot$definition == "Metastatic",]

# Recode gene expression as difference between primary and met
identical(
  annot[annot$definition == "PrimarysolidTumor", "patient"],
  annot[annot$definition == "Metastatic", "patient"]
)
genes.delta <- genes[annot$definition == "PrimarysolidTumor",] -
  genes[annot$definition == "Metastatic",]

dim(genes.delta)
dim(miRNA.delta)

library(exprso)
set.seed(1)
how <- "buildRF"

res.null <- lapply(1:ncol(genes.delta), function(g){

  print(g)

  y.i <- genes.delta[,g]

  library(exprso)
  e.i <- exprso(miRNA.delta, y.i)
  ms <- ctrlModSet(func = "modPermute")
  ss <- ctrlSplitSet(func = "splitSample", percent.include = 67)
  fs <- ctrlFeatureSelect("fsCor")
  gs <- ctrlGridSearch(func = "plGrid", top = 128, how = how, fold = NULL)
  boot.i <- plMonteCarlo(e.i, B = 250, ctrlSS = ss, ctrlFS = fs,
                         ctrlGS = gs, ctrlMS = ms)
  getWeights(boot.i)
})
```

```
# Name and save results
res.df <- lapply(1:length(res.null), function(r){
  data.frame("r" = colnames(genes.delta)[r], res.null[[r]])
})
res.df <- do.call(plyr::rbind.fill, res.df)
write.csv(res.df, file = paste0("5b-DeltaDelta-NULL-", how, "-LOG.csv"))
```

```
setwd("/home/thom/Dropbox/Partners/Thin_Thom/wPaper")
`%+%` <- function(a, b) paste0(a, b)

# CALCULATE p-value FOR RANDOM FOREST RMSE
build <- "buildRF"
null <- read.csv("5b-DeltaDelta-NULL-" %+% build %+% "-LOG.csv", row.names = 1)
actual <- read.csv("5a-DeltaDelta-" %+% build %+% "-LOG.csv", row.names = 1)
pvals <- lapply(unique(null$r), function(i){

  data.frame(
    "id" = i,
    "p" = wilcox.test(null[null$r == i, "valid.rmse"],
                      actual[actual$r == i, "valid.rmse"],
                      alternative = "greater")$p.value,
    "null" = mean(null[null$r == i, "valid.rmse"]),
    "actual" = mean(actual[actual$r == i, "valid.rmse"])
  )
})
pvals <- do.call("rbind", pvals)
pvals.df <- pvals[,"p",drop = FALSE]
rownames(pvals.df) <- pvals[,"id"]
colnames(pvals.df) <- "buildRF"

# VIEW ERROR BASED ON INDEX
view <- pvals[order(pvals$p), "id"][1]
plot(c(actual[actual$r %in% view,"valid.rmse"],
       null[null$r %in% view,"valid.rmse"]),
     col = c(rep("red", 50), rep("blue", 250)))

# FDR-ADJUST p-value
pvals.df$buildRF <- p.adjust(pvals.df$buildRF)
RFsig <- rownames(pvals.df[pvals.df$buildRF < .05,,drop=FALSE])

# RANK NODE PURITY VALUES
sig <- actual
sig[is.na(sig)] <- 0
sig <- sig[sig$r %in% RFsig,]
sig.sub <- sig[,c(1, grep("hsa", colnames(sig)))]
sig.sub[,-1] <- apply(sig.sub[,-1], 1, function(x) rank(x, ties.method =
"min") / length(x))
dim(sig.sub) # OK

# CALCULATE IMPORTANCE
importants <- aggregate(. ~ r, FUN = mean, sig.sub)
rownames(importants) <- importants[,1]
importants <- importants[,-1]
dim(importants) # OK

# GET TOP 2.5% SCORES
allscores <- as.vector(as.matrix(importants))
plot(hist(allscores))
q <- quantile(allscores, .975)

edge <- propr:::wide2long(importants)
edge <- edge[edge$value > q,]
dim(edge) # OK
write.csv(edge, file = "6-Fig4-edges-LOG.csv")
rownames(edge) <- NULL
sink("EdgeList.txt")
knitr::kable(
  edge,
  "latex"
)
sink()
```

```
library(igraph)
g <- graph_from_data_frame(edge[,c("variable", "id")], directed = FALSE)
V <- names(V(g))
sizes <- degree(g)
sizes[grepl("ENSG", names(sizes))] <- 3

library(AnnotationDbi)
library(org.Hs.eg.db)
names <- select(org.Hs.eg.db, keys = names(sizes),
                keytype = "ENSEMBL", columns = "SYMBOL")
names[names$ENSEMBL == "ENSG00000232679", "SYMBOL"] <- "LINC01705"
i <- grepl("hsa", names$ENSEMBL)
names[i, "SYMBOL"] <- gsub("hsa\\.(mir\\.)?", "", names$ENSEMBL)[i]

# SAVE RESULTS
table1 <- pvals
table1$padj <- p.adjust(pvals$p)
table1 <- table1[table1$padj < .05,]
table1 <- merge(table1, names, by.x = 1, by.y = 1)
table1$lfc <- abs((table1$actual - table1$null) / table1$actual * 100)
table1[,c("null", "actual", "lfc", "p", "padj")] <- round(table1[,c("null",
"actual", "lfc", "p", "padj")], 4)
final <- table1[order(table1$padj), c("id", "SYMBOL", "null", "actual", "lfc",
"p", "padj")]
rownames(final) <- NULL
options(scipen=999)
sink("Table1.txt")
knitr::kable(
  final,
  "latex"
)
sink()

# Customize layout
l <- layout_with_kk(g)
i <- grep("hsa.mir.4444.2", V)
l[i,] <- l[i,] + c(-.5,.75)
i <- grep("hsa.mir.1296", V)
l[i,] <- l[i,] + c(0,-.75)
i <- grep("hsa.mir.548o.2", V)
l[i,] <- l[i,] + c(1,0)
i <- grep("hsa.mir.423", V)
l[i,] <- l[i,] + c(-.4,0)
i <- grep("hsa.mir.4484", V)
l[i,] <- l[i,] + c(-1.5,0)
i <- grep("hsa.mir.1296", V)
l[i,] <- l[i,] + c(-.5,0)
i <- grep("hsa.mir.3684", V)
l[i,] <- l[i,] + c(0,-.2)

jpeg("6-Fig4-network-LOG.jpg",
     width = 10, height = 10, units = "in", res = 600)
plot(g,
     vertex.shape = ifelse(grepl("hsa", V), "circle", "square"),
     vertex.color = ifelse(grepl("hsa", V), adjustcolor("lightblue", alpha.f
= .25),
                           adjustcolor("salmon", alpha.f = .25)),
     vertex.size = sizes * 2,
     vertex.frame.color=NA,
     vertex.label = ifelse(sizes >= 3, names$SYMBOL, NA),
     vertex.label.color = "black",
     vertex.label.cex = .7,
     vertex.label.family = "sans",
```

```
     layout = l)
dev.off()
```

```
setwd("~/Dropbox/Thin_Thom/wPaper")
library(devtools)
library(tidyverse)
# install_github("tpq/miSciTools")
library(miSciTools)
library(annotables)

genes <- read_csv("6-Fig4-edges-LOG.csv") %>%
  left_join(grch38, by = c("id" = "ensgene")) %>%
  distinct(symbol) %>%
  pull(symbol)

universe <- read_csv("2-DESeq2-genes.csv") %>%
  filter(padj < 0.05) %>%
  left_join(grch38, by = c("X1" = "ensgene")) %>%
  pull(symbol)

annot <- read_tsv("gsea/msigdb/c5.bp.v6.1.symbols.gmt", col_names = F) %>%
  rename(pathway = X1) %>%
  select(-X2)


aa <- annot[, 1:2]
colnames(aa) <- c("pathway", "gene")
for(cc in colnames(annot)[-c(1, 2)]) {
  a <- select(annot, pathway, cc) %>%
    na.omit()
  colnames(a) <- c("pathway", "gene")
  aa <- bind_rows(aa, a)
}
aa <- arrange(aa, pathway)

sum(genes %in% aa$gene)

gsea <- simpliGSEA(genes, universe, aa$gene, aa$pathway)
q <- p.adjust(gsea[[1]])


gsea <- simpliGSEA(genes, universe, kegg$symbol, kegg$path)

genes[genes %in% kegg$symbol]
```
